# Inhibition of SARS-CoV-2 spike protein palmitoylation reduces virus infectivity

**DOI:** 10.1101/2021.10.07.463402

**Authors:** Ahmed A. Ramadan, Karthick Mayilsamy, Andrew R. McGill, Anandita Ghosh, Marc A. Giulianotti, Haley M. Donow, Shyam S. Mohapatra, Subhra Mohapatra, Bala Chandran, Robert J. Deschenes, Arunava Roy

**Author notes:** Address correspondence to Arunava Roy, Department of Molecular Medicine, University of South Florida, 12901 Bruce B. Downs Blvd., MDC 4138, Tampa, FL 33612, USA.

## Abstract

Spike glycoproteins of almost all enveloped viruses are known to undergo post-translational attachment of palmitic acid moieties. The precise role of such palmitoylation of the spike protein in membrane fusion and infection is not completely understood. Here, we report that palmitoylation of the first five cysteine residues of the c-terminal cysteine-rich domain of the SARS-CoV-2 spike are indispensable for infection and palmitoylation deficient spike mutants are defective in trimerization and subsequent membrane fusion. The DHHC9 palmitoyltransferase interacts with and palmitoylates the spike protein in the ER and Golgi, and knockdown of DHHC9 results in reduced fusion and infection of SARS-CoV-2. Two bis-piperazine backbone-based DHHC9 inhibitors inhibit SARS-CoV-2 spike protein palmitoylation and the resulting progeny virion particles released are defective in fusion and infection. This establishes these palmitoyltransferase inhibitors as potential new intervention strategies against SARS-CoV-2.

## Introduction

The last two decades have seen the emergence of three major coronavirus (CoV) outbreaks. The first was the Severe Acute Respiratory Syndrome-CoV (SARS-CoV) in 2002, then Middle East Respiratory Syndrome-CoV (MERS-CoV) in 2012 and most recently, the SARS-CoV-2 virus causing the COVID-19 pandemic. β-Coronaviruses are enveloped, positive-stranded RNA viruses that express a spike protein on the surface giving the virus the appearance of a crown or corona, in Latin. The SARS-CoV-2 spike glycoprotein (S) is a 1273 amino acids type I membrane protein that binds to the ACE2 receptor on the host cell to initiate infection, viral uptake, and cell-cell fusion (1, 2). Structurally, the unprocessed S protein precursor consists of an N-terminal signal sequence for endoplasmic reticulum (ER) insertion and a large ectodomain (ER luminal - virion exterior) composed of glycosylation sites, a receptor binding domain (RBD), a trimerization domain, and two proteolytic cleavage sites (S1/S2 and S2’) that are required for the conformational changes that present the fusion peptide domain for membrane insertion (3, 4). On the cytosolic side of the single membrane spanning domain is a short endodomain that contains a cysteine rich domain (CRD) capable of undergoing S-acylation/palmitoylation (5, 6).

Protein palmitoylation is the reversible posttranslational addition of palmitate, from palmitoyl-CoA, onto the side chain of cysteine residues via a thioester linkage (7). The reaction is catalyzed by a family of palmitoyl-acyltransferases (PATs) containing a consensus DHHC (Asp-His-His-Cys) sequence (8, 9). While originally described as a lipid anchor for peripheral membrane yeast proteins Ras2 and Yck2 (10, 11), palmitoylation is now known to be a common posttranslational modification that occurs on more than 30% of all cellular proteins (12). Palmitoylation regulates membrane association, trafficking, vesical fission and fusion, and protein stability (13, 14). Palmitoylation of viral proteins has been implicated in replication, viral assembly, budding, and cell fusion (15). For example, palmitoylation of murine coronavirus (MHV) spike is essential for virion assembly and infectivity (16). Moreover, a palmitoylation deficient recombinant MHV virus was shown to be deficient in cell fusion and syncytia formation (17). Similar results were observed in SARS-CoV, where mutagenesis of the first five cysteines of the C-terminal CRD leads to a palmitoylation deficient spike protein with reduced syncytia formation ability (18, 19).

In SARS-CoV-2 infected cells, the S protein is synthesized on rough endoplasmic reticulum (RER) where they are co-translationally embedded into the ER membrane via its N-terminal ER retention signal (20). Subsequently, the S protein trimerizes and are trafficked to the ER-Golgi intermediate compartment (ERGIC), the Golgi network and ER derived double membrane vesicles (DVM) where nascent virion particle assembles by budding into these membranous structures. A C-terminal membrane-spanning ER retrieval signal (ERRS) prevents the S protein from being fully released into the lumen of the ER via the secretory pathway (21, 22). During its passage through the secretory pathway, the S protein is folded, post-translationally modified, and cleaved at the S1/S2 cleavage site by furin or furin-like proteases (23, 24). Eventually, membrane-bound carrier proteins transfer the newly-assembled virions to the plasma membrane for egress (25). In addition to this, a fraction of spike traverses the secretory pathway to the plasma membrane, where they bind ACE2 on uninfected cells leading to multinucleated syncytia (23). This allows for cell-cell spread of the virus without the release of virion particles and also serves to shield the virus from immune detection (23). In the case of SARS-CoV S protein and the influenza virus HA protein, palmitoylation of viral glycoproteins has been shown to precede proteolytic cleavage and glycosylation, but not trimerization which occurs in the ER (15). This suggests that palmitoylation of viral glycoproteins occurs in the late ER, ERGIC and the cis-Golgi cisternae, followed by furin cleavage and glycosylation in the trans-Golgi (15). Evidence that this is may hold true for SARS-CoV-2 also, has recently been reported (26).

Protein palmitoylation is carried out by a family of 23 mammalian DHHC proteins (8). However, information on which of these PATs palmitoylate CoV S proteins has been sparse. Pull-down experiments by Gordon et al. identified DHHC5 palmitoyltransferase and its auxiliary protein, Golga7 as interacting with the SARS-CoV-2 spike protein (27). Recent observations made by Wu et al. and others have indicated that DHHC5 is associated with the palmitoylation of SARS-CoV-2 S protein (6, 28, 29). In contrast, Masquita et al. recently reported that SARS-CoV-2 S protein is mainly palmitoylated by DHHC20, but DHHC8 and 9 also play a role (26). A metabolic modeling study by Santos-Beneit et al. designed to identify potential anti-COVID drug targets identified protein palmitoylation as a prominent target (30). These studies provide support to the notion that inhibitors of PATs may be developed as potential anti-viral drugs (31). The unmet challenge to date is to develop high affinity, specific, PAT inhibitors. The most widely used inhibitor, 2-bromopalmitic acid (2-BP) is a non-metabolizable palmitate analog with no detectable preference toward any specific DHHC PAT (32). Moreover, 2-BP is also known to inhibit other lipid metabolizing enzymes (33). We have previously reported more specific PAT inhibitors based on a Bis-piperazine scaffold (34). In this report, we show that these novel compounds inhibit palmitoylation of SARS-CoV-2 spike proteins resulting in reduced virus infectivity.

Here, we present our studies on the role of palmitoylation of the SARS-CoV-2 spike protein using a pseudotyped luciferase lentivirus system as well as SARS-CoV-2 virus infection. We employ a number of CRD cysteine mutants of the spike protein to evaluate the propensity of different cysteine clusters to be palmitoylated. We then evaluate the role of spike palmitoylation on trimerization, S1/S2 furin cleavage, transport through the secretory pathway, release on mature virion particles, binding to the ACE2 receptor, cell-cell syncytia formation and infection. We identify DHHC9 as a PAT palmitoylating the SARS-CoV-2 spike protein and demonstrate its co-localization and physical interaction with the spike protein in transfected cells as well in SARS-CoV-2 infected cells. Finally, we extended our studies to include SARS-CoV-2-mNG (SARS-CoV-2 stably encoding mNeonGreen) and show that PAT inhibitors also reduce SARS-CoV-2 infection and syncytia formation. Together, these results establish DHHC9 as a potential target against SARS-CoV-2 and identify lead compounds for the intervention of the SARS-CoV-2 lifecycle and infectivity.

## Results

### SARS-CoV-2 spike protein is palmitoylated on multiple sites

Cysteines in close proximity to transmembrane helices have higher propensity to be palmitoylated (35). The C-terminal CRD of SARS-CoV-2 spike protein has 10 highly conserved juxtamembrane cysteine residues, 9 of which were predicted to be potential sites of palmitoylation by the palmitoylation prediction server at http://csspalm.biocuckoo.org/online.php (36) (Fig. 1A). To investigate the role of palmitoylation at these 10 Cysteine residues, we grouped them into 4 clusters - C1 (C1235, 1236), C2 (C1240, C1241, 1243), C3 (C1247, 1248, 1250) and C4 (C1253, 1254), and mutated each cluster to serine (Fig. 1A). We also generated a ΔC mutant in which all 10 cysteines were mutated to serine. The Acyl-PEGyl Exchange Gel-Shift (APEGS) Assay was used to assess the palmitoylation of wild-type (WT) and cysteine mutant S proteins, which employs mPEG-maleimide alkylation to label palmitoylated cysteine residues to study the effect of these mutations on the palmitoylation of the S protein. After PEGylation, the samples were separated by SDS–PAGE and visualized by chemiluminescence using a monoclonal antibody specific for the S2 fragment of the Spike protein. The mPEG minus APEGS reactions served as a control and confirmed the observed gel shifted palmitoylated bands were due to PEGylation at the available cysteine residues. The C1, C2, and C4 cysteine clusters reduced palmitoylation significantly, whereas the ΔC mutation led to undetectable levels of palmitoylation (Fig. 1B). Mutating the C3 cluster appears to be less important for palmitoylation (Fig. 1B). The S protein shifted to several higher molecular weight bands representing different populations of the spike protein with varying degrees of palmitoylation (Fig. 1B). GAPDH, which is also known to undergo palmitoylation (37) was used as a positive control for the APEGS reaction and as a loading control to show equal protein loading per well.

**FIGURE 1:**
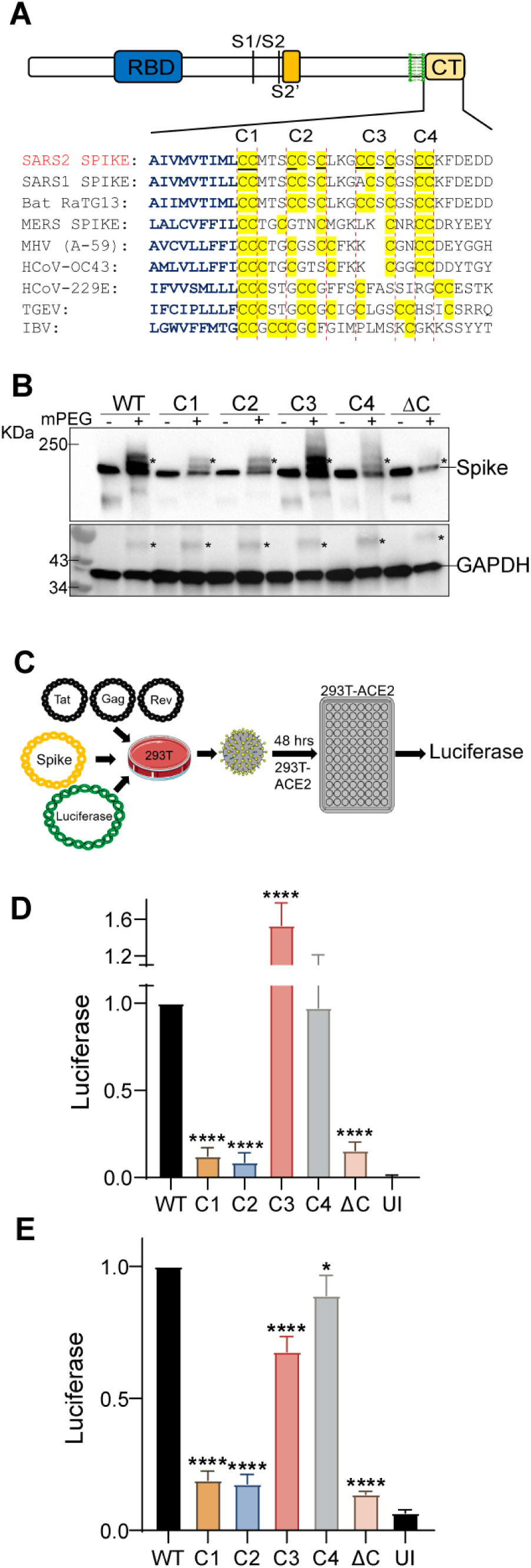
Palmitoylation of SARS-CoV-2 spike protein and effect of different cysteine clusters mutation of the S protein on pseudotyped lentivirus infection. **A.** Sequence alignment of the carboxy-terminal tails of the spike protein from the indicated coronaviruses. The transmembrane region residues are depicted in blue and the conserved cysteine residues in the c-terminal cysteine rich domain (CRD) are highlighted in yellow. The 10 cysteines of the SARS-CoV-2 spike protein (S) CRD were grouped as four clusters – C1 (C1235, 1236), C2 (C1240, C1241, 1243), C3 (C1247, 1248, 1250) and C4 (C1253, 1254). Palmitoylation site prediction algorithm predicted 9 of the 10 underlined cysteine residues as potential site of palmitoylation (http://csspalm.biocuckoo.org/online.php). **B.** Plasmids with WT SARS-CoV-2 spike protein (S) or the indicated cysteine mutants (C1-4 and ΔC, where all 10 cysteines are mutated to serine) were transfected into HEK293T cells and 48 h later, palmitoylation of the spike protein was assessed by the Acyl-PEGyl Exchange Gel-Shift (APEGS) Assay using anti-spike protein antibody. Addition of mPEG results in slower migrating species, indicated by an asterisk (top panel). GAPDH serves as a loading and palmitoylation control (bottom panel). **C.** Schematic of the luciferase reporter SARS-COV-2 spike pseudotyped lentivirus system used. **D.** HEK293T-ACE2 or Caco-2 **(E)** cells were infected with lentivirus pseudotyped with WT spike or its cysteine cluster mutants for 48 h and pseudovirus infection measured by quantifying the luciferase signal. UI represents uninfected control. Data shown are relative to the WT pseudovirus and are averages of the results of at least three independent experiments ± SD. (*p < 0.05; **p < 0.01; ***p < 0.001, ****= p <0.0001 (One-Way ANOVA)).

Next, we employed a luciferase reporter SARS-COV-2 spike pseudotyped lentivirus system to study the effect of these cysteine cluster mutations on the cellular entry and infectivity of the S protein by measuring the luciferase activity 48 h after pseudovirus infection of HEK293T-ACE2 or Caco-2 cells (Fig. 1C). Compared to S WT, the ΔC mutant was highly defective in infection of HEK293T-ACE2 cells (Fig. 1D). Clusters C1 and C2 mutations were also defective in infection to levels similar to the ΔC mutant (Fig. 1D). In contrast, clusters C3 and C4 mutations had almost no diminution of pseudovirus entry. Similar results were observed with Caco-2 cells which expresses ACE-2 endogenously (Fig. 1E). Together, these data indicate that the first five cysteine residues of the SARS-CoV-2 spike protein CRD (C1235, 1236, 1240, 1241 and 1243) are the most functionally important residues to be palmitoylated for infection of ACE-2 expressing cells.

### Mutation of Cluster C2 (C1240, C1241, 1243) of S protein abrogates trimerization

Next, to investigate if the observed reduction in the infectivity of the C1, C2 and ΔC mutant pseudotyped lentiviruses are due to defects in the intracellular processing of the nascent S protein, we tested its trimerization and intracellular furin cleavage at the S1/S2 site by resolving the WT and the mutant S proteins expressed in 293T cells on native-PAGE. Relative band intensities of the three forms of the S protein – S2 fragment, full length, and trimer were normalized to WT being 1.0. As shown in Fig. 2A and B, compared to the WT protein, mutants C2 and ΔC were severely defective in trimer formation. Spike protein trimerization is known to be essential for its membrane fusion and thus, infection ability (38). Furin mediated cleavage at the S1/S2 site of the spike protein is a gain-of-function unique to SARS-CoV-2, which is absent in SARS-CoV and other SARS-related coronaviruses (39). There was no detectable difference in spike furin cleavage between WT and the mutants (Fig. 2A and B).

**FIGURE 2:**
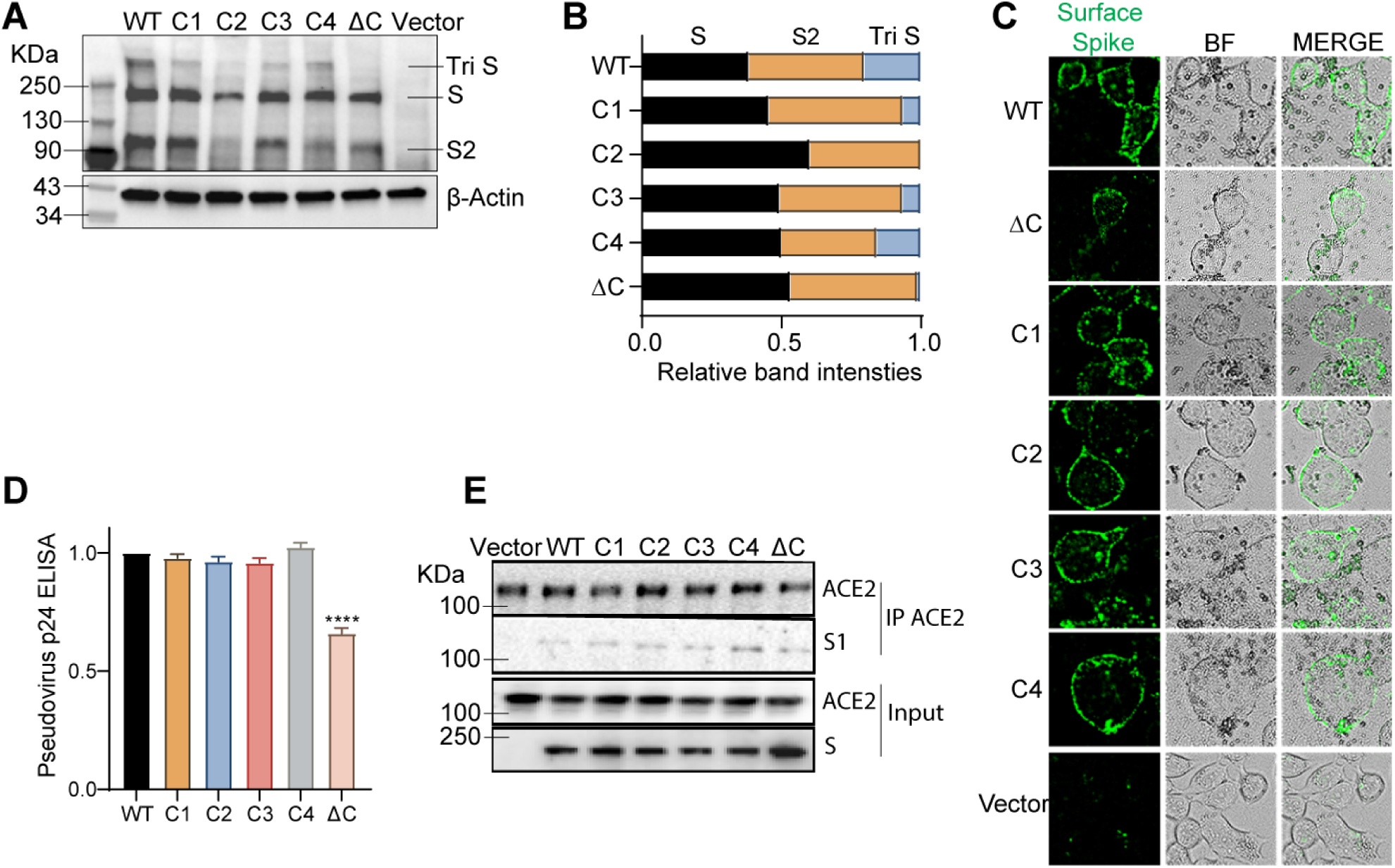
Effect of cysteine cluster mutations of SARS-CoV-2 S protein on its trimerization, plasma membrane localization, egress and ACE2 binding. **A.** Native PAGE showing trimeric spike (Tri S), monomeric spike (S) and the S2 fragment of the WT and different cysteine cluster mutants of the S proteins. Actin is included as loading control. **B.** Relative normalized intensities of tri S, S and S2 bands in panel A. **C.** Surface immunofluorescence assay for the S protein was performed by using anti-S protein antibodies on unpermeabilized HEK293T cells 48 h after transfection with WT or cysteine cluster mutant S proteins. BF indicates bright field. **D.** HEK293T cells were transfected with plasmids for SARS-CoV-2 spike pseudotyped lentivirus and 48 h later, supernatant containing pseudotyped lentivirus particles (WT and cysteine mutants) were assayed for pseudovirus egress by ELISA against the lentivirus (HIV) p24 protein. **E.** Co-immunoprecipitation assay. Plasmids expressing WT or different cysteine cluster mutant S proteins were transfected into HEK293T-ACE2 cells. 48 h after transfection, ACE2 was immunoprecipitated and the presence of S protein was assayed by immunoblot using antibody specific to the S1 subunit. ACE2 immunoprecipitation was also confirmed. Inputs were quantitated by immunoblot (bottom panels).

Lentivirus based pseudotyped virus assembly and budding occurs at the host plasma membrane. Therefore, we investigated the effect of the cysteine cluster mutations on the transport of the S protein to the plasma membrane. For this, we performed surface immunofluorescence assay (SIF) and immunostained the S protein on non-permeabilized cells. We found that only the ΔC mutant exhibited moderate reduction in its surface expression, while the rest of the cysteine cluster mutations had no measurable effect (Fig. 2C). This indicates that the reduced infectivity of the C1 and C2 mutant pseudotyped lentivirus is not due to defects in their ability to be transported to the plasma membrane.

Next, we tested the effect of these mutants on the packaging and egress of the S-pseudotyped lentivirus particles. For this, we collected cell-free S-pseudotyped lentivirus supernatants and compared their egress by performing ELISA against the lentivirus capsid protein, p24. Our results show that none of the cysteine cluster mutants had any effect on the egress of the pseudotyped lentivirus (Fig. 2D). However, the ΔC mutant showed a moderate, but significant reduction, indicating that only mutating all 10 cysteines has a measurable effect on pseudotyped lentivirus egress.

The effect of spike palmitoylation on ACE2 receptor binding was examined by co-immunoprecipitation. ACE2 was immunoreacted with anti-ACE2 antibody followed by detection of spike by immunoblotting with anti-spike antibody. There appeared to be no difference in ACE2 binding comparing WT to the cysteine cluster mutants (Fig. 2E). Together, these experiments establish that ΔC, and to a lesser extent C1 and C2 exhibit reduced trimerization of the spike protein, whereas furin cleavage, cell surface localization, pseudovirus egress, and ACE2 binding remains unaltered.

### Mutating the CRD diminishes S protein mediated membrane fusion and syncytia formation

CoV spike proteins play an indispensable role in the fusion between the viral envelope and the target cell membrane, or between an infected cell and an uninfected cell membrane (40, 41). Both are essential steps in virus infection and entry, and in cell to cell spread and pathogenesis. Therefore, we investigated the effect of the cysteine cluster mutations on the fusogenic potential of the SARS-CoV-2 S protein. For this, we transfected HEK293T-EGFP (HEK293T cells stably transfected with EGFP) with S WT or its mutant plasmids. 293T-ACE2 cells were stained with a red cell tracker dye and 6 h later, the two cells were mixed at 1:1 ratio, and co-cultured together. When the GFP+S expressing cells and the ACE2 expressing red cells fuse and form a syncytium, it will appear as yellow under a fluorescent microscope (Fig. 3A). First, we compared S WT to the ΔC mutant and observed that the palmitoylation deficient spike protein exhibited a marked reduction in syncytia formation (Fig. 3B and C). Subsequently, we tested all the cysteine cluster mutants and, consistent with our pseudovirus infection observations (Fig. 1D and E), we observed that C1 and C2 cluster mutations have the greatest effect on syncytia formation (Fig. 3D and E). C3 and C4 clusters had lower, but significant effects on syncytia formation. These results collectively show that the first 5 cysteines of the SARS-CoV-2 S CRD (C1235, 1236, 1240, 1241, 1243) play the most important role in fusogenic activity.

**FIGURE 3:**
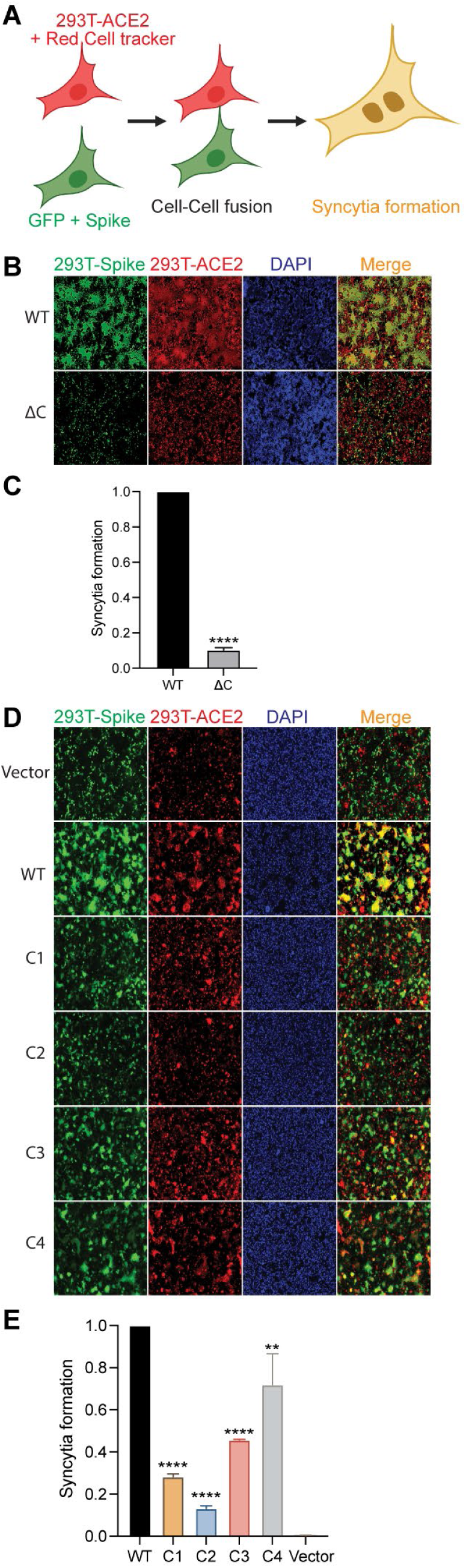
SARS-CoV-2 S protein palmitoylation is required for cell-cell fusion and syncytia formation. **A.** Schematic representation of dual fluorescent syncytia formation assay. HEK293T-EGFP cells were transfected with WT-S or its cysteine mutant plasmids. After 24 h, cells were detached and mixed with HEK293T-ACE2 cells labeled with cell tracker red CMPTX dye. After another 24 h, syncytia formation was evaluated by visualizing the yellow fluorescence formed by the fusion of the green and red cells. **B.** Dual fluorescent syncytia formation assay with WT-S and SΔC and **(C)** quantification of cell-cell fusion ability by measuring the pixel density of the observed syncytia. D-E. Same as in B and C, but, with cysteine cluster mutants of the S protein.

### Reducing DHHC9 levels decreases SARS-CoV-2 spike palmitoylation

Since palmitoylation of the S protein is an indispensable step for CoV infection, we sought to identify the DHHC PAT responsible for palmitoylating the spike CRD. Gordan et.al found that the SARS-CoV-2 spike protein interacts with DHHC5 and Golga7 (27). Golga7 has been previously identified as an accessory protein of DHHC9 and DHHC5 (42, 43). We thus knocked down DHHC5 and DHHC9 individually using siRNA in HEK293T cells and 72h later confirmed their knock down efficiencies using RT-PCR and WB (Fig 4A and B). We found that both enzymes were knocked down efficiently. Interestingly, knock down of DHHC5 led to a 1.8-fold upregulation of DHHC9 (Fig. 4A), whereas knock down of DHHC9 did not affect DHHC5 expression. The change in mRNA levels resulted in commensurate changes in protein levels (Fig. 4B). To examine this further, we measured mRNA levels of a panel of DHHC genes following knock down of DHHC5 and DHHC9. Knock down of DHHC5 resulted in a compensatory upregulation of DHHC9, 15 and 20 (Fig. S1). However, knock down of DHHC9 did not result in any such compensatory upregulation of any other PAT (Fig. S1).

**FIGURE 4:**
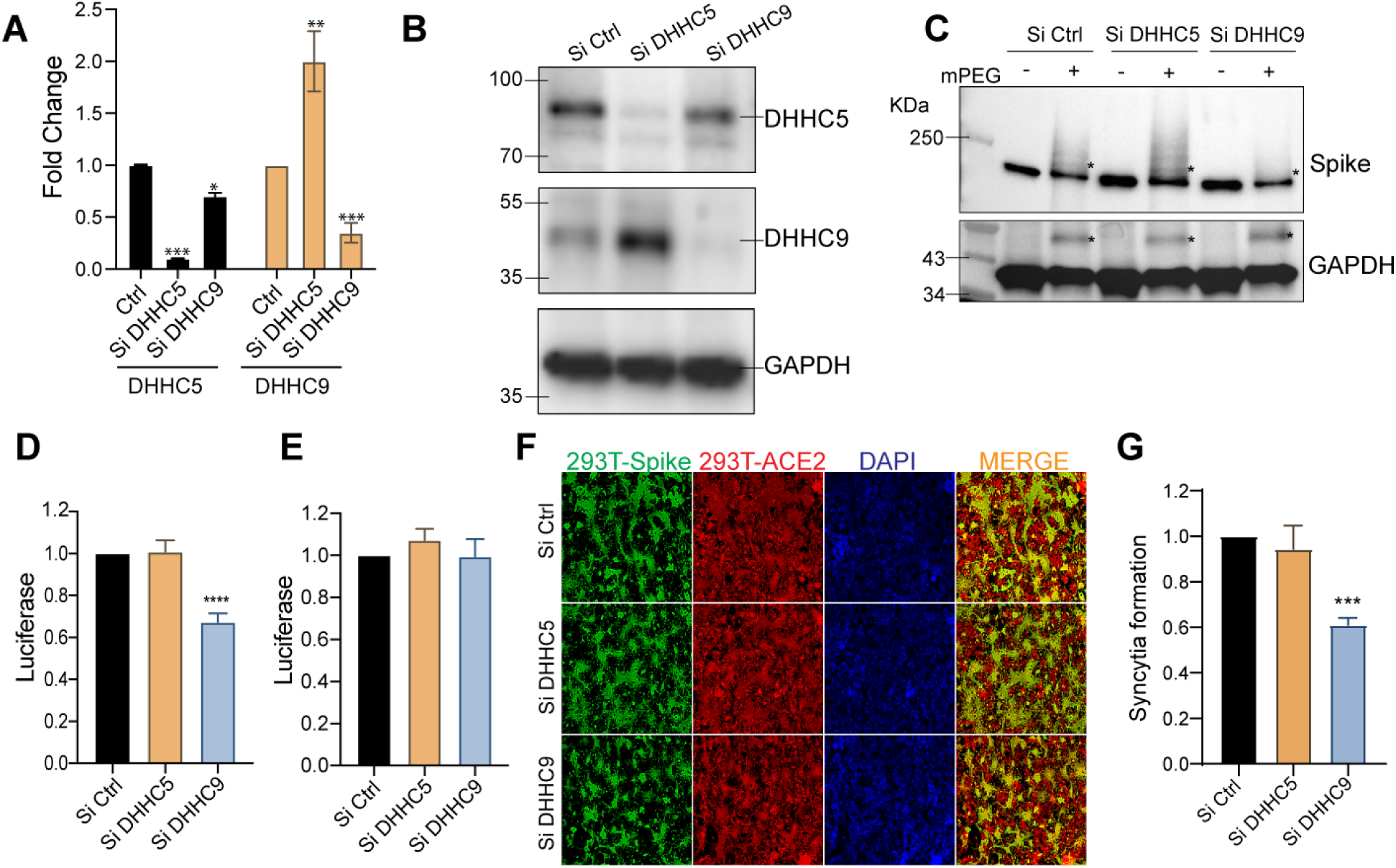
Evaluation of DHHC5 and DHHC9 acyltransferases as SARS-CoV-2 spike palmitoylating enzymes. **A.** DHHC5 and DHHC9 acyltransferases were knocked down using siRNA for 72 h in HEK293T cells and their respective mRNA levels evaluated by RT-PCR. **B.** Immunoblots assessing the efficiency of the siRNA knocked down of DHHC5 and DHHC9. **C.** APEGS assay to evaluate the role of DHHC5 and DHHC9 on S protein palmitoylation in HEK293T cells. **D.** HEK293T-ACE2 cells were infected with WT spike protein pseudotyped lentivirus isolated from HEK293T cells treated with control siRNA or siRNA against DHHC5 or DHHC9. Results are normalized to control siRNA set at 1.0. **E.** HEK293T-ACE2 cells were knocked down for DHHC5 or DHHC9 and 48 h later, infected with WT spike protein pseudotyped lentivirus derived from untreated HEK293T cells. Further 48 h later, pseudovirus infection was measured by quantifying the luciferase signal. Data shown are relative to the control siRNA. **F.** Dual fluorescent syncytia formation assay. HEK293T-EGFP cells were transfected with siRNA targeting DHHC5 or DHHC9 and 48 h later, transfected WT-S plasmids. 24 h later, the cells were detached and mixed with HEK293T-ACE2 cells labeled with cell tracker red CMPTX dye. After another 24 h, syncytia formation was evaluated by visualizing the yellow fluorescence formed by the fusion of the green and red cells. **G.** Quantification of cell-cell fusion ability of the data described in (E) by measuring the pixel density of the observed syncytia. All data shown are averages of the results of at least three independent experiments (3 fields each) ± SD. *=p < 0.05; **=p < 0.01; ***=p < 0.001, ****= p <0.0001 (unpaired t test).

The APEGS assay was used to examine the palmitoylation of the spike protein following knock down or DHHC5 or DHHC9. Reduction of DHHC9 significantly reduced spike palmitoylation (Fig 4C). In contrast, in DHHC5 knocked down cells S protein palmitoylation increased, presumably due to the increase in DHHC9, but we cannot rule out that DHHC15 and DHHC20 may also contribute since they are similarly upregulated in a DHHC5 knockdown (Fig. S1). As expected, palmitoylation of GAPDH was unaffected upon knockdown of both DHHC5 or DHHC9 (44) (Fig 4C, lower pane).

To evaluate the effect of DHHC5 and DHHC9 down regulation on the ability of S protein pseudotyped lentivirus to infect ACE2 bearing cells, we generated lentivirus particles from cells in which DHHC5 and 9 levels have been knock down. Reduction of DHHC9, but not DHHC5, resulted in a significant reduction in infection of HEK293T-ACE2 cells (Fig. 4D). In contrast, knock down DHHC9 or DHHC5 in the HEK293T-ACE2 cells had no effect on the infection by pseudovirus derived from untreated cells (Fig. 4E). Thus, S protein palmitoylation is required for infection of HEK293T-ACE2 recipient cells, but not for any downstream event following infection. We next tested if syncytia formation requires DHHC5 or DHHC9. As seen in Fig. 4F and G, knock down of DHHC9, but not DHHC5, resulted in significantly reduced syncytial formation.

### DHHC9 co-localizes and interacts with the SARS-CoV-2 spike protein both in transfected and infected cells

To determine whether DHHC9 interacts with the SARS-CoV-2 spike protein, we performed a Co-IP experiment with FLAG tagged DHHC9 or its catalytic mutant, DHHA9 Mut, in HEK293T cells co-transfected with the spike protein. DHHC9 and the spike protein physically interact and the interaction was not dependent on the palmitoyltransferase activity of DHHC9 (Fig. 5A). In contrast, a similar Co-IP experiment with Myc-tagged Golga7 failed to detect an interaction with the spike protein (Fig. 5B).

**FIGURE 5:**
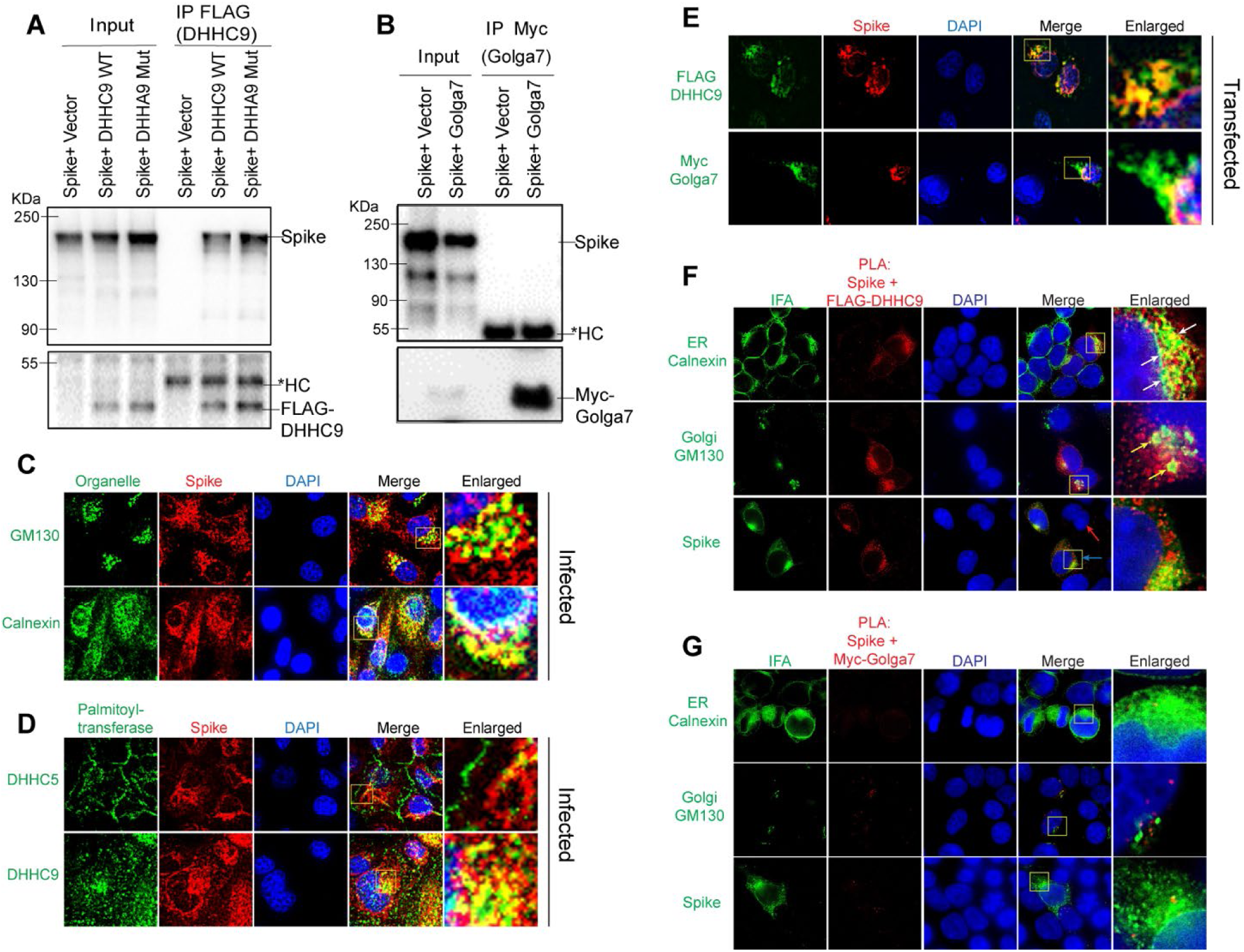
Interaction of DHHC9 with SARS-CoV-2 spike protein. **A.** Immunoprecipitation of FLAG tagged DHHC9 with spike protein. HEK293T cells were transfected with either FLAG-DHHC9, or its catalytic inactive mutant of DHHC9 (FLAG-DHHA9 Mut) and an untagged spike protein and 48 h later, immunoprecipitated with FLAG antibodies. The respective empty vectors were used as controls. HC, heavy chain of IgG. **B.** Immunoprecipitation of Myc tagged Golga7 with spike protein. HEK293T cells were transfected with Myc tagged Golga7 and an untagged spike protein and 48 h later, immunoprecipitated with anti-Myc antibodies. **C.** Co-localization of spike protein with the cis-Golgi marker, GM130 or the endoplasmic reticulum marker, Calnexin. Vero-E6-ACE2 cells were infected with SARS-CoV2 (MOI 0.01) and 48 h later, immunostained for the indicated proteins. **D.** Similar experiment as in (B) showing co-localization between spike protein and endogenous DHHC5 and DHHC9. **E.** HEK293T cells were transfected with either untagged spike protein with FLAG-DHHC9 or Myc-Golga7 and 48 h later, immunostained with anti-FLAG or anti-Myc antibodies together with anti-spike antibodies, to show co-localization of transfected spike with exogenous DHHC9 and Golga7. **F.** Proximity ligation assay (PLA) to detect the physical proximity between the spike protein and FLAG-DHHC9. Experiment was performed as in (D), and PLA was performed with the indicated antibodies followed by immunofluorescence (IFA) against either GM130 or Calnexin to visualize the sub-cellular localization of the detected PLA spots. **G.** PLA to detect the physical proximity between the spike protein and Myc-Golga7. Experiment same as in F.

Next, to examine whether the spike protein interacts with DHHC9 in infected cells, we performed immunofluorescence experiments in Vero-E6-ACE2 cells infected with SARS-CoV-2 for 48 h. Co-localization (yellow) with the endoplasmic reticulum marker, Calnexin and the cis-Golgi marker, GM130 indicates that the S protein localizes to both the ER and the Golgi apparatus (Fig. 5C). Similar observations were made in HEK293T cells transfected with FLAG-DHHC9, Myc-Golga7 or the spike protein, where all of these proteins were found to localize to the ER and the Golgi network (Fig. S2A). Following this, we performed co-localization experiments between the spike protein and DHHC5 and 9 in SARS-CoV-2 infected Caco-2 cells (Fig. 5D). In agreement with previous observations (44, 45), we observed DHHC5 to be localized predominantly on the cell surface, but in addition, some DHHC5 could be observed at other intracellular locations (Fig. 5D). Because the S protein is primarily localized to the ER and the Golgi, we did not observe significant co-localization between S and DHHC5 (Fig. 5D, upper panel). However, under similar conditions, DHHC9 extensively co-localized with spike protein (Fig. 5D, lower panel). To further confirm the spike protein’s co-localization with DHHC9 and to exclude the possible non-specific staining due to the DHHC9 antibody, we performed similar co-localization experiments in HEK293T cells transfected with FLAG DHHC9 and Myc-tagged Golga7. Significant co-localization was observed between DHHC9 and the spike protein (Fig. 5E, top panel). Myc Golga7, also co-localized with the spike protein, albeit to a lesser degree (Fig. 5E, bottom panel).

The proximity ligation assay (PLA) provides a better method to assess direct interactions (<40nm) between proteins in cells (46). When PLA was performed, we observed robust interactions between the spike protein and FLAG DHHC9 (Fig. 5F). We also performed immunofluorescence against Calnexin and GM130 in the same experiment and found that the PLA signal between the S protein and DHHC9 localizes partially to the Golgi apparatus, but almost entirely to the ER network (Fig. 5F, arrows). This suggests that palmitoylation of spike occurs primarily in the ER and secondarily in the ERGIC, as Calnexin is also found in the ERGIC (47, 48). In a similar experiment, where PLA was performed between the spike protein and Golga7, we failed to detect a significant PLA signal, indicating again that spike and Golga7 do not directly interact with each other (Fig. 5G). The specificity of the PLA signal was confirmed by isotype IgG controlled PLA reaction which did not produce any signal (Fig. S2B).

### SARS-CoV-2 infection and syncytia formation in Caco-2 cells requires DHHC9 dependent palmitoylation of spike protein

Having established that DHHC9 plays a major role in the palmitoylation of the SARS-CoV-2 spike protein in-vitro using pseudovirus, we directed our attention to study the SARS-CoV-2 virus. SARS-CoV-2-mNG, a fluorescently labeled virus, is based on the 2019-nCoV/USA_WA1/2020 strain isolated from the first reported SARS-CoV-2 case in the US (49). SARS-CoV-2-mNG is a recombinant virus in which the mNeonGreen gene has been inserted into ORF7 of the viral genome. This recombinant virus exhibits similar plaque morphology, viral RNA profile, and replication kinetics compared to the original clinical isolate (49). Thus, this virus can be used to determine the efficiency of SARS-CoV-2 infection and syncytia formation. Caco-2 cells transfected with DHHC9 or DHHC5 siRNA were infected with SARS-CoV-2-mNG and 24, 48 and 72 h post-infection, monitored for mNeonGreen expression. At 24h post-infection, the mNeonGreen signal was not measurably reduced after knockdown of DHHC5 or DHHC9 (Fig. 6A, B and S3). However, at 48 and 72 h post-infection, the mNeonGreen signal in the DHHC9 knocked down cells reduced significantly. We reason that, as the cells were infected with SARS-CoV-2-mNG harvested from WT Vero-E6 cells, the S protein on these virus particles were efficiently palmitoylated and knock down of the respective PATs in the recipient Caco-2 cells had no effect on the infection process and the expression of mNeonGreen at 24 h post infection. However, with time (48 and 72 h), nascent virion particles increasingly harbored palmitoylation-deficient spike protein which resulted in reduced mNeonGreen signal and lower levels of infection of neighboring cells. The size of the syncytia was also reduced in the DHHC9 knocked down cells compared to the control knocked down cells at 72 h post-infection (Fig. 6A). There was no measurable change in syncytia formation after knock down of DHHC5, suggesting that DHHC9 plays a key role in the palmitoylation of the spike protein during SARS-CoV-2 infection.

**FIGURE 6:**
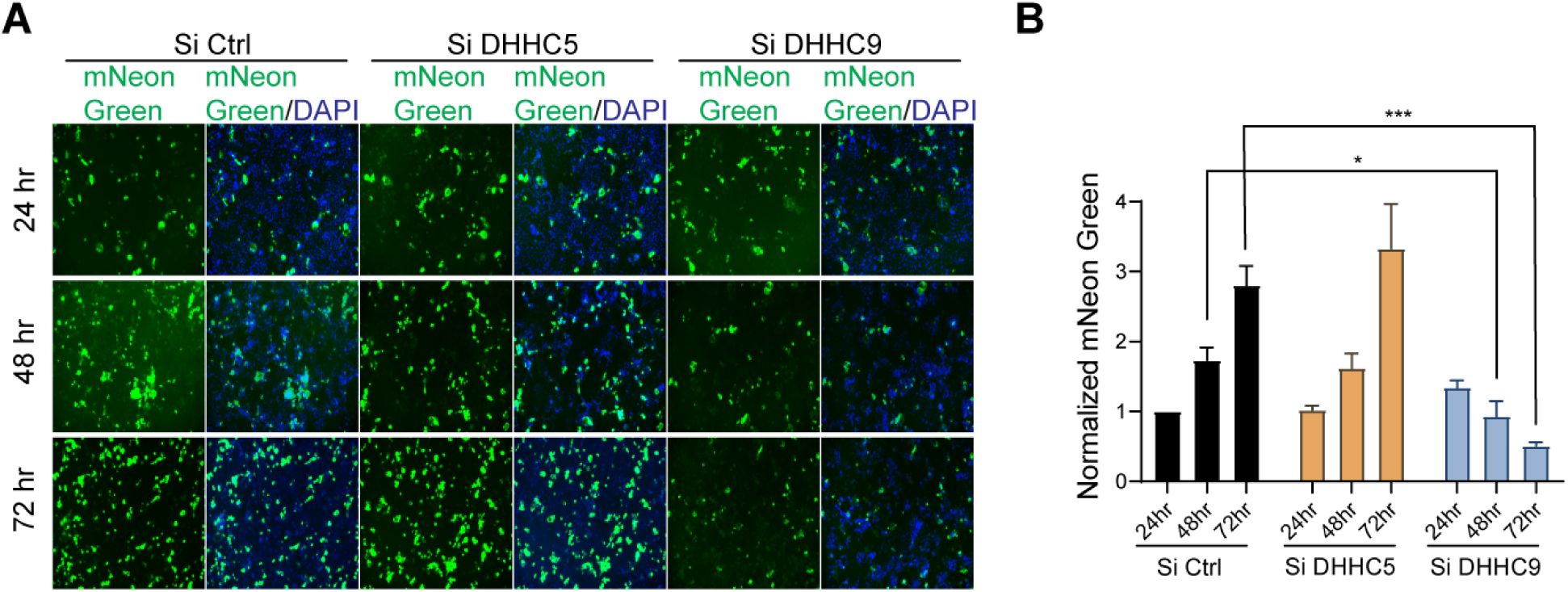
Effect of DHHC9 knockdown on SARS-CoV-2 infection in Caco-2 cells. **A.** Caco-2 cells were knocked down for DHHC5 or DHHC9 and 48 h later, infected with icSARS-CoV-2-mNG (SARSCoV-2 stably encoding mNeonGreen; MOI 0.1). 24, 48 and 72 h post-infection, the cells were fixed, nucleus stained with DAPI and visualized under a fluorescence microscope. **B.** mNeonGreen signal from (A) was quantitated, normalized to DAPI and plotted to show the effect of the respective acyltransferase knocked down on icSARS-CoV-2-mNG infection. All data shown are averages of the results of at least three independent experiments (3 fields each) ± SD. *=p < 0.05; **=p < 0.01; ***=p < 0.001, ****= p <0.0001 (unpaired t test).

### DHHC9 inhibitors inhibit SARS-CoV-2 spike palmitoylation, fusogenicity and infectivity

Having identified DHHC9 as a SARS-CoV-2 spike protein palmitoylating enzyme, we investigated whether inhibiting DHHC9 would inhibit SARS-CoV-2 infection. The availability of validated and specific PAT inhibitors are fairly limited (50). 2-bromopalmitate (2-BP), the most widely used PAT inhibitor promiscuously inhibit a wide range of enzyme utilizing active site cysteine residues (32, 51–53). Previously, using a scaffold ranking approach to screen for novel inhibitors of the yeast homolog of DHHC9, members of our group identified a number of bis-piperazine backbone based compounds (34). Two leads from this study, compounds 13 and 25, inhibited palmitoylation at low micromolar concentrations. Compound 13 has the lead functional group, 2-(3,5-bis-trifluoromethylphenyl)-ethyl, at positions R1 and R3, while compound 25 has 4-tert-butyl-cyclohexyl-methyl at the R1 position and 2-(3,5-bis-trifluoromethyl-phenyl)-ethyl at position R3 (Fig. 7A). We evaluated the palmitoylation inhibitory potential of compounds 13 and 25 against the SARS-CoV-2 spike protein.

**FIGURE 7:**
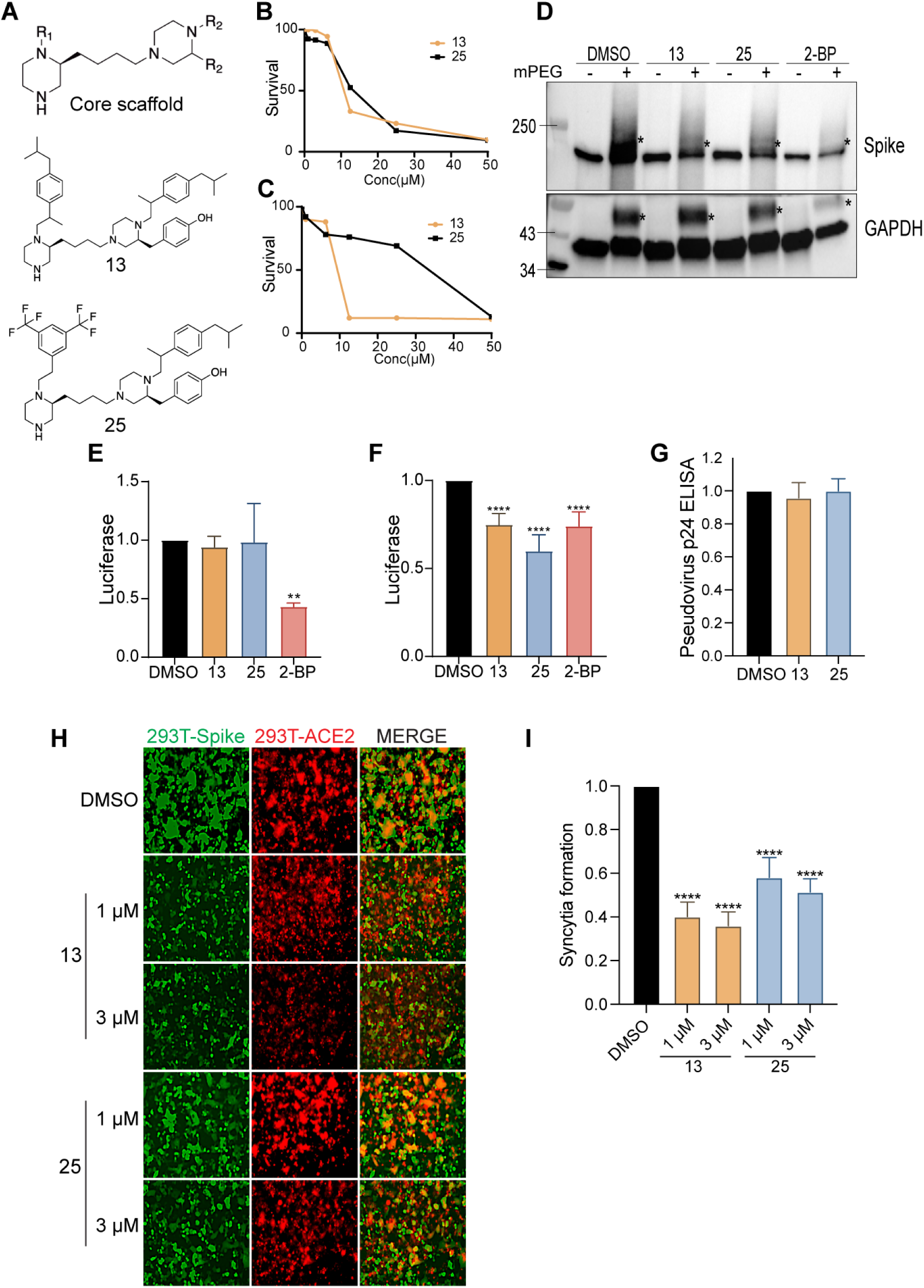
Inhibition of SARS-CoV-2 infectivity by DHHC9 inhibitors. **A.** Chemical structures of compounds 13 and 25. **B** and C. XTT cell viability assay to access the toxicity of compounds 13 and 25 on HEK293T and Caco-2 cells respectively. Survival is plotted relative to DMSO, vehicle only control. **D.** APEGS Assay to evaluate the palmitoylation of the WT-S protein after treatment with compounds 13 and 25. HEK293T cells were treated with the maximal non-toxic concentration of each compound (3µM) or the broad spectrum, non-specific acyltransferase inhibitor 2-BP (10µM) for 12 h before transfection with the spike plasmid. 48 h later, APEGS assay was performed. DHHC9 independent palmitoylation of GAPDH is included as a control. **E.** HEK293T-ACE2 cells were pretreated with either compounds 13 (3µM) or 25 (3µM) or 2-BP (10µM) for 12 h and then infected with luciferase reporter lentivirus pseudotyped with WT-S protein. Luciferase activity was measured 48 h later. **F.** HEK293T cells were pretreated with either compounds 13 (3µM), 25 (3µM) or 2-BP (10µM) for 12 h and then transfected with plasmids required to produce luciferase reporter lentivirus pseudotyped with WT-S protein. The pseudovirus collected was used to infect HEK293T-ACE2 cells and luciferase activity was measured after 48 h. **G.** HEK293T cells were pretreated with compounds 3 (3µM) or 25 (3µM) or 2-BP (10µM) and then transfected with plasmids required to produce lentivirus pseudotyped with WT-spike protein. 48 h later, the supernatant containing pseudotyped lentivirus particles were assayed for pseudovirus egress by ELISA against the HIV p24 protein. **H.** Dual fluorescent syncytia formation assay. HEK293T-EGFP cells were pretreated with the indicated concentrations of compounds 13 and 25 for 12 h and then transfected with WT S plasmid. 24 h later, the cells were detached and mixed with HEK293T-ACE2 cells labeled with cell tracker red CMPTX dye. Syncytia formation was evaluated at 24 h by visualizing the yellow fluorescence formed by the fusion of the green and red cells. **I.** Quantification of cell-cell fusion ability by measuring the pixel density of the observed syncytia in (H). All data shown are averages of the results of at least three independent experiments ± SD. *=p < 0.05; **=p < 0.01; ***=p < 0.001, ****= p <0.0001 (unpaired t test).

The compounds were first tested for toxicity on HEK293T and Caco-2 cells and found to have no observable toxicity at concentrations below 3µM (Fig. 7B and C, respectively). We next examined whether compounds 13 and 25 inhibited the SARS-CoV-2 spike protein palmitoylation using APEGS assay. Compared to the vehicle control, compounds 13 and 25 (3µM) reduced palmitoylation of the spike protein in HEK293T cells by 37% and 44% respectively (Fig. 7D). The non-specific PAT inhibitor, 2-BP, also inhibited spike palmitoylation, but required 10µM concentration. In the same experiment, we found that GAPDH palmitoylation is reduced by the non-specific inhibitor, 2-BP, but not by compounds 13 and 25 (Fig. 7D), consistent with the specificity previously observed for these compounds (34).

We next examined the effect of these palmitoyltransferase inhibitors on the cellular entry and infectivity of SARS-COV-2 spike pseudotyped lentivirus on HEK293T-ACE2 cells. First, we treated the recipient HEK293T-ACE2 with inhibitors and found that there was no reduction in luciferase signal upon treatment with compounds 13 and 25 (Fig. 7E). However, 2-BP caused a significant reduction in pseudovirus infection in this experiment, implying that it interferes with lentivirus endocytosis (Fig. 7E). This also signifies that compounds 13 and 25 do not affect any step downstream of infection. In a second series of experiments, HEK293T cells were treated with compounds 13, 25 or 2-BP and pseudovirus produced from these cells were tested for their ability to infect untreated HEK293T-ACE2 cells. In this case, we observed a significant reduction in the luciferase signal (Fig. 7F) indicating that inhibition of spike palmitoylation resulted in a pseudovirus with reduced ability to infect cells. Under similar conditions, 2-BP also reduced luciferase signal significantly. We also found that compounds 13 and 25 do not reduce the quantity of pseudovirus released from the producer cells, indicating that there is no measurable effect on lentivirus packaging and egress (Fig. 7G). Furthermore, we observed that inhibition of spike palmitoylation by compounds 13 and 25 reduces its fusogenicity in a syncytia formation assay. Treatment with compound 13 (1μM and 3μM) resulted in a 58% and 60% reduction in syncytia formation, respectively. Compound 25 (1μM and 3μM) similarly reduced syncytia formation by 45% and 50%, respectively (Fig. 7H and I).

### Compounds 13 and 25 inhibit SARS-CoV-2 infection in cell culture

We tested the effect of these palmitoyltransferase inhibitors on SARS-CoV-2 infection using the SARS-CoV-2-mNG. Caco-2 cells pretreated with compounds 13, 25, or 2-BP and infected with SARS-CoV-2-mNG exhibited dose dependent reduction in mNeonGreen signal after 72 h of infection (Fig. 8A, B and S4). The size of the syncytia were also reduced in the inhibitor treated cells compared to the control vehicle treated cells (Fig. 8A). This indicates that compounds 13 and 25 effectively inhibits SARS-CoV-2 spike protein palmitoylation causing a reduction in the infection competent virus released to subsequently infect neighboring cells.

**FIGURE 8:**
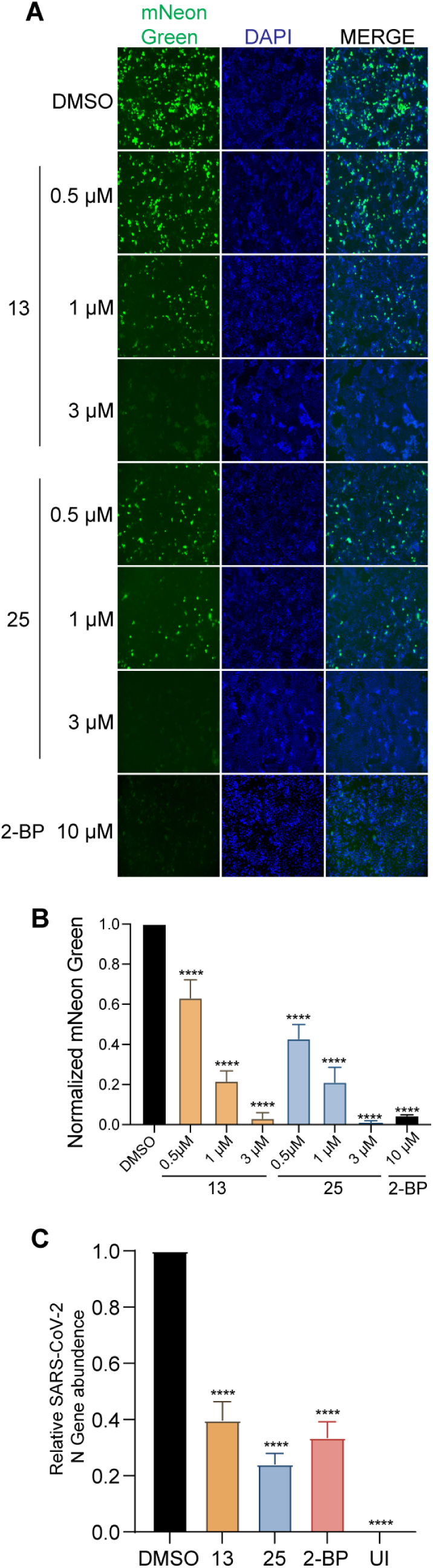
Compounds 13 and 25 inhibits SARS-CoV-2 infection. **A.** Caco-2 cells were pretreated with the indicated concentrations of compounds 3 and 25 for 12 h and then infected with icSARS-CoV-2-mNG (MOI 0.1). Post-infection, the cells were continued to be incubated in the presence of the respective compound dilutions. 72 h later, the cells were fixed, nucleus stained with DAPI and visualized under a fluorescence microscope. 10µM 2-BP was also included in this experiment. **B.** mNeonGreen signal from (J) was quantitated (6 fields from each experiment), normalized to DAPI and plotted to show the effect of the respective concentrations of the DHHC9 PAT inhibitors on icSARS-CoV-2-mNG infection. Data shown are averages of the results of at least three independent experiments ± SD (One-way ANOVA) **C.** Caco-2 cells were pretreated with compounds 13 (3µM), 25 (3µM) or 2-BP (10µM) for 12 h and then infected with SARS-CoV-2 (MOI 0.1). Post-infection, the cells were continued to be incubated in the presence of the respective compound dilutions. 72 h later, the virus containing supernatants were collected and used to infect Vero-E6 ACE2 cells. 24 h after this infection, the Vero-E6 ACE2 cells were collected, RNA extracted and the SARS-CoV-2 N gene quantified using RT-qPCR. UI represents uninfected cells. Data shown are averages of the results of at least three independent experiments ± SD (unpaired t test). *=p < 0.05; **=p < 0.01; ***=p < 0.001, ****= p <0.0001.

Next, virus containing supernatants were collected 72 h after SARS-CoV-2 infection of Caco-2 cells pre-treated with the inhibitors and used to infect Vero-E6 ACE2 cells. After 24 h, infection was quantitated by measuring the appearance of the SARS-CoV-2 N gene by real-time RT-PCR. SARS-CoV-2 virus isolated from Caco-2 cells treated with compounds 13 and 25 resulted in a 60% and 76% reduction viral infection of Vero-E6 cells (Fig. 8C).

## Discussion

In this study, we demonstrate that the SARS-CoV-2 spike protein is palmitoylated on a cluster of conserved cysteines residues on the cytosolic domains of the SARS-CoV-2 spike protein. Mutating all 10 cysteine residues to serine (ΔC) eliminates all detectable palmitoylation, but mutating individual clusters of cysteines suggests that not all cysteine residues are palmitoylated equivalently. For example, mutating clusters C1 (C1235, C1236), C2 (C1240, C1241, C1243), and C4 (C1253, C1254) significantly reduces, but does not eliminate spike palmitoylation. In contrast, spike palmitoylation is unaffected by mutating the C3 (C1248, C1249, C1250) cluster. Individual cysteine clusters are also functionally different. Mutating C1 and C2 clusters reduce infection of ACE-2 expressing cells to the same extent as ΔC, indicating that palmitoylation of the juxtamembrane cysteines are the most important for infection. Mutating the C3 cluster does not reduce palmitoylation or influence infection. Interestingly, mutating the C4 cluster reduces overall palmitoylation, but does not reduce infection suggesting that palmitoylation of the C4 cysteines may have other roles in the viral life cycle.

Palmitoylation of SARS-CoV-2 spike occurs in the ER and Golgi and results in partitioning into detergent resistant, cholesterol and sphingolipid rich membrane microdomains (26). In contrast, palmitoylation defective spike is localized in detergent soluble membrane fractions and results in 35% less spike on the cell surface. We also find that the ΔC mutant of the spike protein decreases surface expression by 40% and taken together, these results show that palmitoylation is important, but not essential for the surface expression of the spike protein. The spike protein appears to be able to access alternate trafficking routes depending on palmitoylation. We have also shown that palmitoylation is not required for ACE2 receptor binding. However, despite being capable of binding ACE2 and a modest decrease in membrane expression, virus harboring non-palmitoylated spike fail to infect host cells. One reason could be that membrane fusion depends on surface density of spike proteins (54). Palmitoylation dependent clustering of the spike protein in infected cell lipid rafts could increase spike density on the membrane to support membrane fusion. Alternatively, palmitoylation of spike may play a more direct role in membrane fusion.

Since the palmitoylation sites of the SARS-CoV spike protein CRDs are in the vicinity of the membrane bilayer, it is likely that the cytoplasmic tail folds back creating a membrane anchor. The potential to palmitoylate up to 10 cysteine residues would create a strong membrane anchor for the endodomains of the spike protein. While it is possible that changes in the spike protein endodomain via palmitoylation or mutation of the cysteine residues might destabilize the spike protein resulting in its proteasomal clearance, we do not think this is the case. First, we find that the wild-type spike protein and its palmitoylation defective mutants are expressed at approximately the same levels, and, traffic to the plasma membrane and bind to ACE2 at the same levels. However, we observed that the ΔC and C2 cysteine mutants results in a decrease in spike protein trimerization. Previous work with the murine coronavirus spike protein has suggested that trimerization may be important for fusion activity (55) and that the spike protein CRD plays an important role in its oligomerization (6, 56). Pre-fusion, the spike protein typically exists in a metastable conformation. Once it interacts with the host ACE2 receptor, extensive structural rearrangement of the S protein occurs, allowing the virus to fuse with the host cell membrane (40, 57). This S2 domain directed fusion event requires a concerted cooperation between the different domains of the spike trimers (58). It is therefore possible that conformational changes mediated by palmitoylation in the cytoplasmic endodomain of the spike protein, impact the ability of the extracellular ectodomain to adopt the proper conformation required for efficient cell fusion, implying a co-operation between the ecto and endodomains. Alternatively, palmitoylation might simply concentrate the spike protein in membrane microdomains, thus facilitating trimer formation. In addition, extensive palmitoylation of the CRD could also be required to create a stable anchor during the membrane fusion process. One or more of these mechanisms likely results in the failure of non-palmitoylated spike proteins to carry out efficient membrane fusion.

S-acylation of viral proteins requires the host cell palmitoylation machinery that, depending on the cell type, consists of up to 23 individual DHHC PAT genes. Identification of the DHHC protein or proteins that palmitoylate the SARS-CoV-2 spike has begun to emerge from several lines of investigation. First, a comprehensive interactome study uncovered an interaction between the SARS-CoV-2 spike protein and Golga7, an auxiliary protein for DHHC5, DHHC9, and possible additional DHHC proteins (27). All animal reservoirs of coronaviruses express both DHHC9 and DHHC20 (59), and observations from the knockdown and overexpression of DHHC8, 9, and 20 further implicate these PATs in the palmitoylation of SARS-CoV-2 spike protein (26). We used several approaches to show that DHHC9 plays a major role in palmitoylating the SARS-CoV-2 spike protein. Co-localization and proximity ligation studies confirm that both, the spike protein and DHHC9, occupy the same spatial compartments inside the cell and they physically interacts with each other in the ER, Golgi and the ERGIC. Knockdown of DHHC9 resulted in a decrease in spike protein palmitoylation, pseudovirus fusion, syncytia formation, and a 55% and 80% reduction of SARS-CoV-2 infection at 48 h and 72 h post inoculation, respectively. Surprisingly, we observed that knockdown of DHHC5 resulted in an increase in palmitoylation of the spike protein. On further investigation, we found that knockdown of DHHC5 resulted in compensatory upregulation of DHHC9, DHHC15 and DHHC20 PATs. This suggests that in addition to DHHC9, DHHC15 or DHHC20 can also palmitoylate the spike protein. Others have also shown that DHHC20 is capable of palmitoylating the SARS-CoV-2 spike protein (5, 26, 60). Among these reports, while Mesquita et al. (26) and Li et al. (60) also found DHHC9 as a potential spike palmitoylating PAT, Puthenveetil et al. did not identify DHHC9 in their screen (5). In this context, it is reasonable to assume that spike protein palmitoylation may be carried out by different DHHC proteins in different cellular compartments during the viral life cycle.

Our work and that of several other groups have led to the suggestion that inhibitors of palmitoylation may be developed into a new class of antivirals (27, 30, 31). To do so will require identification of high affinity, specific inhibitors of DHHC PATs. This has proved to be difficult. Commonly used inhibitors such as 2-bromopalmitte show little specificity, hitting a wide range of thiol containing enzymes (32). We previously developed a high throughput palmitoylation assay and used it to screen a chemical library consisting of 68 unique scaffolds and 30 million unique structures for inhibition of the yeast ortholog of DHHC9 (34). Two compounds based on a bis-piperazine backbone (compounds 13 and 25) were selected to be tested for inhibition of SARS-CoV-2 spike palmitoylation and viral infection. We found that both compounds decreased spike protein palmitoylation, pseudovirus infection and syncytia formation. Further, experiments on SARS-CoV-2 infection established the robust inhibitory effect of compounds 13 and 25 on progeny virion formation. Interestingly, the amino acid sequence of the core domain of DHHC9 is highly homologous to that of DHHC20 (59). Therefore, it will be interesting to evaluate in future studies if compounds 13 and 25 inhibit DHHC20 in addition to DHHC9. Because the spike protein of viruses with zoonotic potential are often found to be palmitoylated by a similar set of PATs (59), compounds 13 and 25 may also have potential anti-viral effect on other existing or emerging viruses.

## Materials and Methods

### Plasmids and HIV-1 derived pseudovirus generation

Plasmids used in this study for the production of pseudo-typed virus were a gift from Jesse D. Bloom’s lab (BEI Resources # NR52516, NR52517, NR52518 and NR52519). The HDM-IDTSpike-fixK (BEI catalog number NR-52514) plasmid expressing a codon-optimized spike from SARS-CoV-2 strain Wuhan-Hu-1 (Genbank NC_045512) under a CMV promoter, was used as the wild-type spike and we performed all the mutagenesis on this codon-optimized spike clone. Third Generation Lenti-virus EGFP plasmid, pLJM1-EGFP, was a gift from David Sabatini (Addgene #19319). 3^rd^ generation lentivirus were produced using the flowing packaging plasmids, which were a gift from Didier Trono’s lab, pMDL (Addgene #12251), pRev (Addgene #12253) and pMD2.G (Addgene #12259) (61).

HIV-1 derived virus particles pseudotyped with full length wild-type and mutant SARS-CoV-2 spike protein were generated by transfecting HEK293T cells as previously described (62). Briefly, plasmids expressing the HIV-1 gag and pol (pHDM-Hgpm2), HIV-1 rev (pRC-CMV-rev1b), HIV-1 Tat (pHDM-tat1b), the SARS CoV2 spike (pHDM-SARS-CoV-2 Spike) and a luciferase/ZsGreen reporter (pHAGE-CMV-Luc2-IRES-ZsGreen-W) were co-transfected into HEK293T cells at a 1:1:1:1.6:4.6 ratio using CalPhos mammalian transfection kit (TaKaRa Clontech, Mountain View, CA #631312) according to manufacturer’s instructions. Fresh media was added after 18 hours and 60 hours later, culture supernatant was collected, clarified by passing through 0.45 um filter and used freshly.

### Cell culture

Human female embryonic kidney HEK293T cells (ATCC) were grown in Dulbecco modified Eagle’s medium (DMEM) supplemented with 10% fetal bovine serum (FBS) (Sigma, #F4135) and Penicillin-Streptomycin (Gibco, #15140148). HEK293T cells constitutively expressing human ACE2 (HEK293T-ACE2 cells) obtained from BEI Resources (#NR52511) were grown in the same media supplemented with hygromycin (100 μg/ml). HEK293T cells constitutively expressing EGFP (HEK293T-EGFP) were established by transducing HEK293T cells with the pLJM1-EGFP containing lentivirus. Lentivirus was produced by transfecting 10^6^ HEK293T cells on a 60 mm plate with pMDL, pRev and pVSVG using CalPhos mammalian transfection kit (TaKaRa Clontech). Next day, fresh media was added, then two days after transfection, the supernatant was harvested, and filtered 0.45-μm filter to remove cell debris. The filtered supernatant containing lentiviral vectors was used to transduce HEK293T cells seeded one day before. 48 hours later, the cells were selected in DMEM supplemented with 2 μg/ml puromycin for one week. The puromycin-resistant population of HEK293T-EGFP cells was found to constitutively express EGFP by fluorescent microscopy. These HEK293T-EGFP cells were used as donor cells in syncytium formation assays. Caco-2 (human colon epithelial cells) cells, obtained from ATCC (HTB-37), were grown in Minimum Essential Medium (Gibco # 11095080.) media supplemented with 20% fetal bovine serum (Sigma, #F4135), Penicillin-Streptomycin (Gibco, #15140148), 1x non-essential amino acid solution (Cytiva, SH3023801) and 10 mM sodium pyruvate (Gibco, # 11360070). All cell lines were incubated at 37°C in the presence of 5% CO_2_.

### CoV spike protein sequence alignment

Amino acid sequences of the S protein used in the alignment were obtained from UniProKB. The accession numbers are SARS-CoV-2 (P0DTC2), SARS-CoV-1 (P59594), Bat RaTG13 (A0A6B9WHD3), MERS (K9N5Q8), MHV [A-59] (P11224), HCoV-OC43 (P36334), HCoV-229E (P15423), TGEV (P07946) and IBV (P11223). Alignment of these sequences was done using Clustal Omega (https://www.ebi.ac.uk/Tools/msa/clustalo/).

### Site-directed mutagenesis

All mutagenesis was done using Q5 Site-directed Mutagenesis Kit Protocol (NEB #E0554S) according to the manufacturer’s instructions using primers in table T1. All mutagenesis was confirmed with sequencing (GENEWIZ, South Plainfield, NJ).

### Transfection

For pseudo-typed virus production, HEK293T cells were transfected using the CalPhos mammalian transfection kit (TaKaRa Clontech) according to the manufacturer’s instructions. For the syncytia formation assay, HEK293T-GFP cells were transiently transfected with wild-type and mutant spike plasmids using TransIT-X2 transfection reagent (Mirus #MIR 6000) according to the manufacturer’s instructions, and fresh media was added after 6 hours.

### SARS-CoV-2 spike pseudotyped virus entry

The assay was done as previously described (62) with minor modification. Briefly, 96-Flat bottom well plates were coated with poly-D-lysine (Gibco, #A3890401) and seeded with 1.25x10^4^ HEK293T-ACE2 or Caco-2 cells per each well. After 12 h, pseudotyped virus with wild-type and mutant spike (with no polybrene) were used for infection. After 48 hours, Steady-Glo Reagent (Promega #E2620) equal to the volume of culture medium in each well was added as per manufacturer’s instructions, cells were allowed to lyse for 5 minutes and then luminescence measured with a microtiter plate reader (Biotek, Winooski, VT).

### Pseudovirus egress

In order to quantitate HIV based pseudovirus egress from HEK293T cells, we performed ELISA against HIV p24 (TaKaRa Clontech #632200) according to manufacturer’s instructions. Briefly, pseudotyped virus containing cell culture supernatant was collected, diluted 1:20, ELISA performed and absorbance measured at 450 nm using a microtiter plate reader (Biotek, Winooski, VT).

### Immunoblotting

Whole cell lysates were prepared using RIPA Lysis Buffer (Thermo scientific #89900) supplemented with a protease inhibitor cocktail (Roche# 11836170001) for 30 min on ice and then sonicated three times at an amplitude setting of 50 with pulses of 15s on and 15s off on a Qsonica Q700 sonicator. The lysates were clarified by centrifugation at 13,000 X g for 15 min at 4°C. Protein concentrations were estimated using Pierce BCA protein assay kit (Thermo scientific #23225) as per manufactural instructions and equal concentration of proteins resolved appropriate SDS PAGE gels. SDS polyacrylamide gels were transferred on Nitrocelluose membranes (GE) by Wet transfer BioRad System at 300 Amps for 90 min at 4°C, membranes were blocked with 5% non-fat milk for 1 hour at room temperature (RT) and were incubated with primary antibodies diluted in PBST solution at 4°C overnight. All primary antibodies used in this study are listed in table T3. Membranes were washed with PBST for 5 minutes, 3 times and subsequently incubated with appropriate secondary antibodies. Blots were then washed three times with PBST for 5 minutes each wash. The immunoreactive bands were developed using Super Signal West Pico chemiluminescent substrate (Thermo scientific #34078) or Super Signal West Femto chemiluminescent substrate (Thermo scientific #34095) depending on the signal strength. Blots were developed on a Bio-Rad ChemiDoc XRS+ System.

### Co-immunoprecipitation (Co-IP)

Whole cell lysates were prepared using Pierce IP Lysis Buffer (Pierce #87788) supplemented with a protease inhibitor cocktail for 30 min on ice and then sonicated three times at an amplitude setting of 30 with pulses of 15s on and 15s off on a Qsonica Q700 sonicator at 4°C. The lysates were then clarified by centrifugation at 13,000 X g for 15 minutes at 4°C and protein concentrations estimated by BCA reaction. For IP, 300 µg of the prepared lysate was incubated with the appropriate antibody (3 µg) and pulled down using protein A Sepharose 6MB (GE healthcare # 17-0469-01). All primary antibodies used in this study are listed in table T3. The immunoprecipitates were washed three times with lysis buffer at 4°C and resolved on SDS-PAGE followed by immunoblotting. Wherever mentioned, light-chain-specific secondary antibodies were used to avoid heavy chain bands in WB of co-IP experiments.

### Acyl-PEGyl Exchange Gel-Shift (APEGS) Assay

To assess the level of protein S-palmitoylation on the SARS-CoV2 spike protein we did APEGS as described previously (63). In brief, HEK293T cells transfected with appropriate plasmids were lysed with the following buffer, 4% SDS, 5 mM EDTA, in triethanolamine buffer (TEA) pH 7.3 with protease inhibitors and PMSF (5 mM). After centrifugation at 20,000 × g for 15 min, proteins in the supernatant were reduced with 25 mM tris(2-carboxyethyl)phosphine (TCEP, Thermo Scientific, #20490) for 1 hour at RT, and free cysteine residues were blocked with 20 mM N-ethyl maleimide (NEM, Sigma #E3876) for 3 hours at RT. To terminate the NEM reaction and wash any residual NEM, pre-chilled methanol: chloroform: H2O (4:1.5:3) was added to the reaction. This wash was repeated three times. Next, the proteins were re-suspended in TEA with 4% SDS and 5 mM EDTA, then incubated in buffer containing 0.2% Triton X-100, 5 mM EDTA, 1 M NH_2_OH, pH 7.0 for 1 hour at RT to cleave palmitoylation thioester bonds. The reaction was terminated as above, and proteins re-suspended in TEA with 4% SDS were PEGylated with 1.33 mM mPEGs (10 kDa, Sunbright, #ME050) for 2 hours at RT to label palmitoylation sites. The reaction was terminated as above; proteins were re-suspended with TEA buffer with 4% SDS. Protein concentration was measured by BCA protein assay. Thereafter, SDS PAGE sample buffer was added and samples were heated at 70 °C for 10 minutes and run on a 7.5% gel for Spike and 12% for GAPDH.

### Syncytium formation assay

HEK293T-EGFP cells (donor cell) were transiently transfected with wild-type and mutant spike plasmids using TransIT-X2 transfection reagent (Mirus #MIR 6000). HEK293T-ACE2 and HEK293T (as a negative control) were stained with CellTracker Red CMTPX Dye (Invitrogen #C34552) according to manufacturer’s instructions. 6 h after transfection, the HEK293T-ACE2 cells were treated with accutase cell detachment reagent and added to the HEK293T-EGFP cells transfected with spike protein, at a 1:1 ratio. 48h after transfection, cells were fixed and imaged using Keyence BX700 fluorescent microscope. Images were quantified using ImageJ.

### Compound Synthesis and Characterization

Bis-piperidines 13 and 25 were synthesized using solid-phase chemistry, purified by RP-HPLC, and characterized by LCMS and 1H NMR as previously described (Hamel L. D., et al. 2016). The compounds were additionally recharacterized by LCMS before performing all biological assays. For LCMS analysis a Shimadzu 2010 LCMS system, consisting of a LC-20AD binary solvent pumps, a DGU-20A degasser unit, a CTO-20A column oven, and a SIL-20A HT auto sampler. A Shimadzu SPD-M20A diode array detector was used for detections. A full spectra range of 190-800 nm was obtained during analysis. Chromatographic separations were obtained using a Phenomenex Gemini NX-C18 analytical column (5 μm, 50 x 4.6 mm ID). The column was protected by a Phenomenex Gemini-NX C18 column SecurityGuard (5 μm, 4 x 3.0 mm ID). All equipment was controlled and integrated by Shimadzu LCMS solutions software version 3. Mobile phases for LCMS analysis were HPLC grade or LCMS grade obtained from Sigma Aldrich and Fisher Scientific. The mobile phases consisted of a mixture of LCMS grade acetonitrile and water (both with 0.1% formic acid for a pH of 2.7). The initial setting for analysis was at 5% acetonitrile (v/v), linearly increasing to 95% acetonitrile over 6 minutes. The gradient was then held at 95% acetonitrile for 2 minutes until linearly decreasing to 5% over 1 minute. From there, the gradient was held until stop for an additional 3 minutes. The total run time was equal to 12 minutes. The total flow rate was set to 0.5 mL/minute. The column oven and flow cell temperature for the diode array detector were set at 40°C. The auto sampler temperature was held at 15°C. 10 μl was injected for analysis.

4-(((2S)-1-(2-(4-isobutylphenyl) propyl)-4-(4-((2S)-1-(2-(4-isobutylphenyl) propyl) piperazin-2-yl) butyl) piperazin-2-yl) methyl) phenol (Compound 13) OC1=CC=C(C[C@@H]2N(CC(C3=CC=C(CC(C)C)C=C3)C)CCN(CCCC[C@@H]4N(C C(C5=CC=C(CC(C)C)C=C5)C)CCNC4)C2)C=C1 LCMS (ESI+) Calculated exact mass for C45H68N4O: 680.54 found [M+H]+:681.55. Retention Time: 4.78 minutes. 90.1% purity by 214 nM.

4-(((2S)-4-(4-((S)-1-(3,5-bis(trifluoromethyl)phenethyl) piperazin-2-yl) butyl)-1-(2-(4-isobutylphenyl) propyl) piperazin-2-yl) methyl) phenol (Compound 25) OC1=CC=C(C[C@@H]2N(CC(C3=CC=C(CC(C)C)C=C3)C)CCN(CCCC[C@@H]4N(C CC5=CC(C(F)(F)F)=CC(C(F)(F)F)=C5)CCNC4)C2)C=C1 LCMS (ESI+) Calculated exact mass for C42H56F6N4O: 746.44, found [M+H]+:747.40. Retention Time: 4.722 minutes. 90.4% purity by 214 nM.

### Cellular toxicity assay

The XTT Cell Proliferation Kit II (Roche # 11465015001) was used for assessing cellular toxicity following manufacturer’s instruction. DMSO was used as a vehicle for all compounds and appropriate dilution of DMSO only was used as a control.

### Gene Silencing

Genes were silenced using siRNAs obtained from IDT, DHHC5 (IDTDNA #290128941) and DHHC9 (IDTDNA #290128950). For each gene, we used a combination of three siRNAs. HEK293T cells were seeded on 24 well plate 24 h before transfection. Transfection of these siRNAs was done using TransIT-X2 (Mirus #MIR 6000) according to the manufacturer’s instructions. Silencing efficacy was checked using qPCR using SYBR Green real-time reagents (Invitrogen # 4367659) and immunoblotting.

### RNA extraction and RT-qPCR

Total RNA was isolated using the RNeasy minikit (Qiagen #74106) following manufacturer’s instructions. On-column DNase digestion was performed by using an RNase-free DNase set (Qiagen #79254). The extracted RNA concentration was estimated using a NanoDrop spectrophotometer (Thermo Scientific), and 1 μg RNA was reverse transcribed by using the High-Capacity cDNA reverse transcription kit (Applied Biosystems #4368814) with random primers, according to the manufacturer’s instructions. For real-time quantitative reverse transcription-PCR (qRT-PCR), the synthesized cDNA was diluted 1:20 and used as a template with Power SYBR Green PCR Master Mix (Applied Biosystems #4367659) on an ABI Prism 7500 detection system (Applied Biosystems). All RNA levels were normalized to β-actin mRNA levels and calculated as the delta-delta threshold cycle (ΔΔC_T_). Primers used in this study are listed in table T2.

### Immunofluorescence assay (IFA) and Proximity Ligation Assay (PLA)

Cells were grown on eight chamber glass slides, and treated as described in the results. After appropriate incubations, the cells were fixed using 4% paraformaldehyde for 15 min, and permeabilized with 0.2% Triton X-100 in PBS for 20 min. The slides were then washed, blocked with Image-iT FX signal enhancer (Invitrogen #I36933) for 30 minutes at 37°C and incubated with primary antibodies for 1 h at 37°C. All primary antibodies used in this study are listed in table T3. After this, the slides were washed three times in PBS and incubated with corresponding fluorescent dye-conjugated secondary antibodies for 30 minutes at 37°C. PLA was performed according to the manufacturer’s instructions using the following kits and reagents: Duolink *in-situ* PLA Probe Anti-Rabbit PLUS (Sigma-Aldrich # DUO92002), Duolink *in-situ* PLA Probe Anti-Mouse MINUS (Sigma-Aldrich # DUO92004), Duolink *in-situ* Detection Reagents Red (Sigma-Aldrich # DUO92008). After completion of IFA or PLA, slides were mounted using mounting medium containing DAPI and observed either by a Keyence BZ-X fluorescence microscope.

### Surface Immunofluorescence Assay

To detect the amount of spike protein localizing to the surface of the cell, HEK293T cells were seeded in 8-chamber glass bottomed slides and transfected with appropriate plasmids. Cells were washed and treated with freshly prepared 0.1% paraformaldehyde and incubated for 10 minutes at 4°C. Thereafter, the cells were washed two times with DMEM with 5% FBS and incubated with anti-spike antibody diluted (1:200) in DMEM with 5% FBS and 0.1% Sodium Azide for 1 h at 37°C. Next, cells were washed four times with DMEM with 5% FBS at 4°C for 10 minutes and incubated with appropriate secondary antibody (1:500) for 30 minutes at 4°C. Finally, the cells were washed, anti-fade reagent added and imaged using a Keyence BX700 microscope.

### SARS-CoV-2 virus stock preparation and titration with plaque-based assays

All replication competent SARS-CoV-2 experiments were performed in a biosafety level 3 laboratory (BSL-3) at the University of South Florida. All viral stocks were produced and isolated from supernatants of Vero-E6 ACE2 cells, cultured in T175 culture flasks to a confluency of 80-90%, and infected with an original passage 2 (P2) SARS-CoV-2 or SARS-CoV-2-mNG (SARS-CoV-2 stably encoding mNeonGreen) virus, at MOI of 0.1 for 72 h, in 10 ml MEM supplemented with 5% FBS. SARS-CoV-2 was obtained from BEI Resources (NR52281), while, SARS-CoV-2-mNG was a kind gift from Dr. PEI-Yong Shi from the University of Texas Medical Branch, Galveston, TX, USA (49). Supernatants were harvested, cleared of cell debris by centrifugation (500g, 10 min) and filtration (0.45 μm), mixed with 10% SPG buffer (ATCC #MD9692), aliquoted and stored at -80°C. Viral titers were quantified by determining the number of individual plaques forming units after 72 h of infection on confluent Vero-E6-ACE2 expressing cells. In brief, viral stocks were serially diluted (10-fold) in serum-free medium and inoculated on 1x 10^5^ Vero E6-ACE2 cells in triplicates in a 48 well plate.

### SARS-CoV-2 infection

All SARS-CoV-2 infections were performed using the same passage 3 SARS-CoV-2 or SARS-CoV-2-mNG virus stocks. Caco-2 cells seeded to a confluency of 70 to 80%, were washed twice in warm serum-free medium and inoculated with the indicated MOI of the appropriate virus, diluted in serum-free medium (5 ml for T75; 2 ml for T25; 1 ml for 6-well plates). Two hours after inoculation cells were washed with complete medium and infection was allowed to proceed for the indicated time points in DMEM supplemented with 2.5% FBS. After infection, media with respective drugs were added and incubated for 72 h. Images were quantified using ImageJ.

### Drug treatments

For drug treatment, cells were treated with indicated concentrations of 2-BP (Sigma, #238422) or compounds 13 and 25 dissolved in DMSO, 12 h prior to infection. Post-infection, cells were continued to be incubated in presence of the respective drugs for the indicated time.

### Statistical analysis and reproducibility

Statistical analysis was performed using GraphPad Prism 9 (San Diego, CA). For two groups, means were compared by two-tailed unpaired Student’s t test. For multiple groups, analysis was done by one-way ANOVA with Dunnett correction for multiple comparisons. P value of < 0.05 was considered statistically significant. Specific statistical test results are indicated in each figure: *= p < 0.05; **= p < 0.01; ***= p < 0.001; ****= p <0.0001.

## Acknowledgements

Funding: This work was supported in part by Public Health Service grant R21 NS090160 to RJD, Public Health Service grant R01 CA180758 to BC, Veterans Affairs Merit Review grant BX005490 and Research Career Scientist Awards IK6BX004212 and IK6BX003778 to SM and SSM.

We gratefully thank the following sources for reagents: The SARS-Related Coronavirus 2, Wuhan-Hu-1 Spike-Pseudotyped Lentiviral Kit was obtained through BEI Resources, NIAID, NIH: SARS-Related Coronavirus 2, Wuhan-Hu-1 Spike-Pseudotyped Lentiviral Kit, NR-52948. Vector pHDM Containing the SARS-Related Coronavirus 2, Wuhan-Hu-1 Spike Glycoprotein was obtained through BEI Resources, NIAID, NIH: Vector pHDM Containing the SARS-Related Coronavirus 2, Wuhan-Hu-1 Spike Glycoprotein, NR-52514. SARS-CoV-2 was deposited by the Centers for Disease Control and Prevention and obtained through BEI Resources, NIAID, NIH: SARS-Related Coronavirus 2, Isolate USA-WA1/2020, NR-52281. SARS-CoV-2-mNG was a kind gift from Dr. PEI-Yong Shi from the University of Texas Medical Branch, Galveston, TX, USA.

---

**Table T1.**
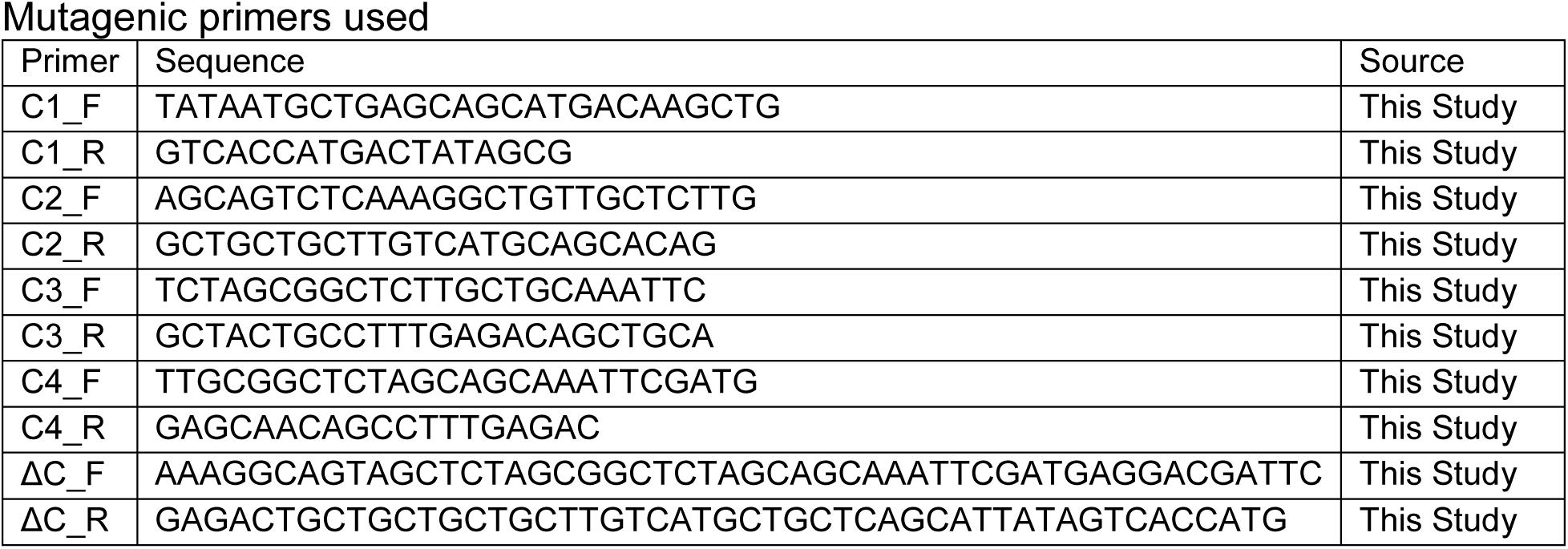

**Table T2.**
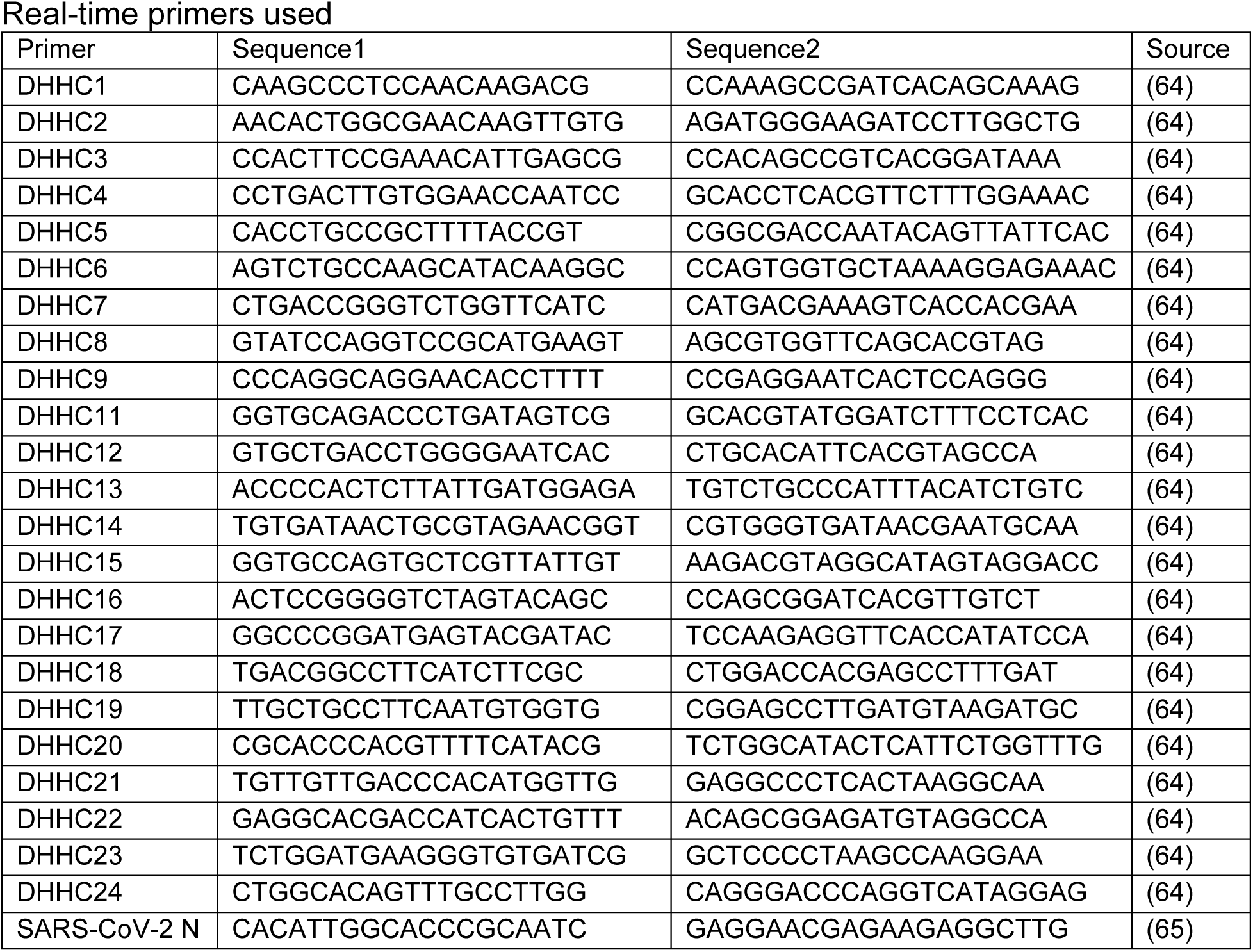

**Table T3.**

Primary antibodies used
| **Antibody** | **Source** | **Catalog Number** |
| --- | --- | --- |
| Spike S2 | GeneTex | GTX632604 |
| Spike S1 | Sino Biological | 40591-MM42 |
| ACE2 | Novus | NBP2-67692 |
| GAPDH | Santa Cruz | sc-47724 |
| Calnexin | Novus | NB300-518 |
| Calnexin | Cell Signaling | 2679P |
| FLAG | Sigma-Aldrich | F1804 |
| FLAG | Sigma-Aldrich | F7425 |
| MYC | Cell Signaling | 71D10 |
| MYC | Cell Signaling | 9B11 |
| DHHC9 | Sigma-Aldrich | HPA031814 |
| DHHC5 | Sigma-Aldrich | HPA014670 |
| β-Actin | Protein-tech | HRP-60008 |
| GM130 | Invitrogen | PA5-95727 |
| GM130 | R&D | AF8199 |

**Fig S1:**
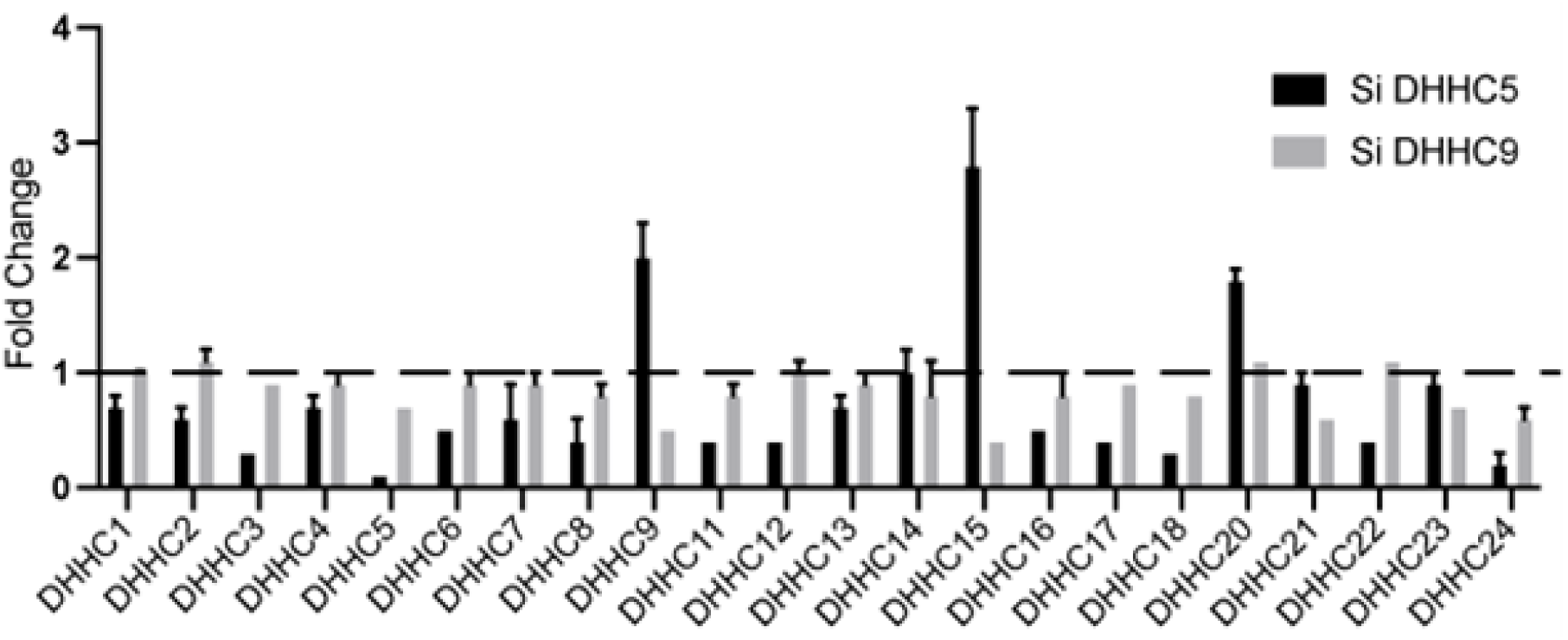
DHHC5 and DHHC9 acyltransferases were knocked down using siRNA for 72 h in HEK293T cells and the indicated acyltransferase mRNA levels were evaluated by RT-PCR.

**Fig S2:**
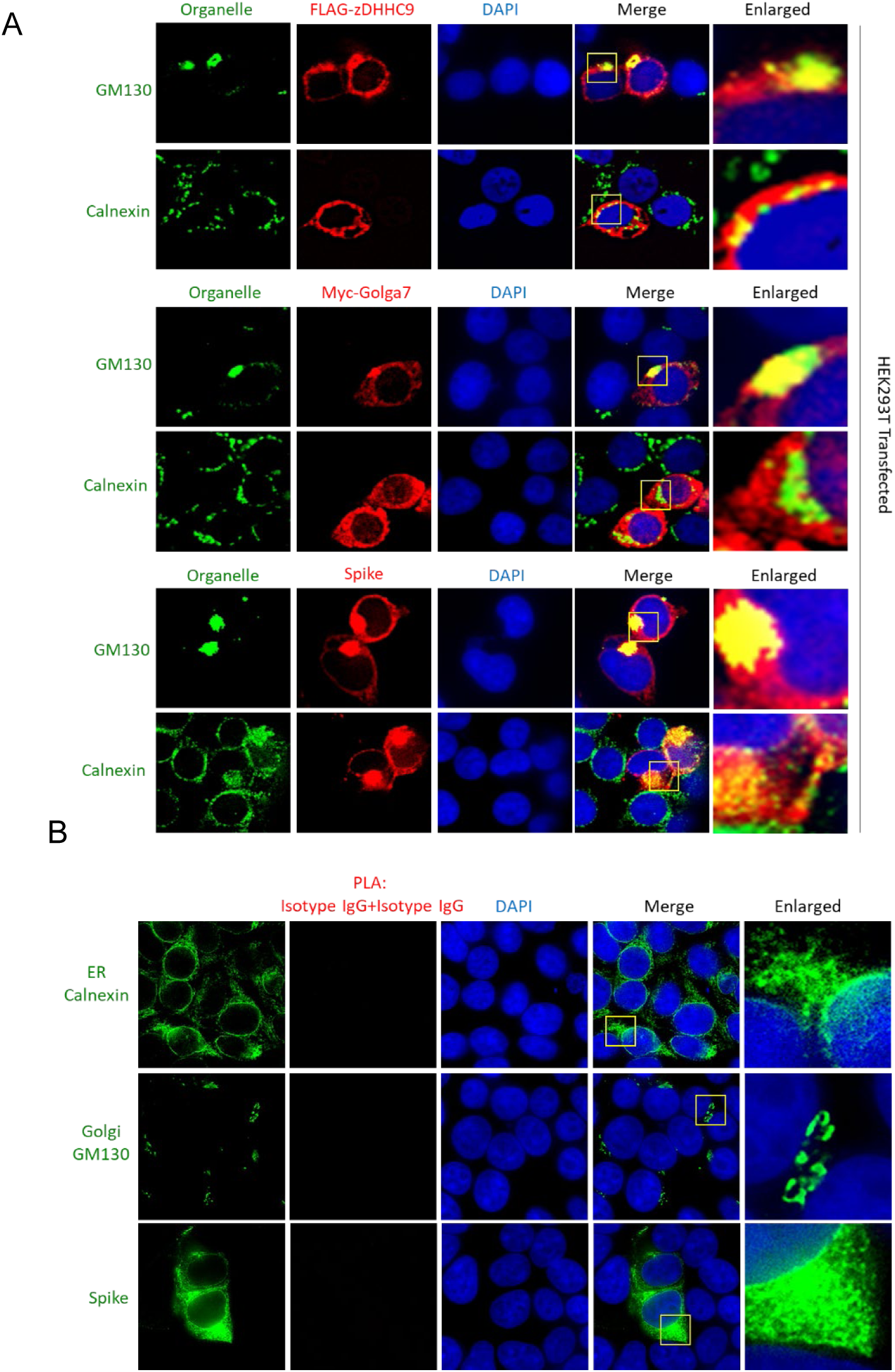
**A.**Co-localization of FLAG-DHHC9, Myc-Golga7 and spike protein with the cis-Golgi marker, GM130 or the endoplasmic reticulum marker, Calnexin. HEK293T2 cells were transfected with the indicated plasmids and 72 h later, immunostained for the indicated proteins. **B.** Control PLA with isotype IgGs showing absence of non-specific PLA signal.

**Fig S3:**
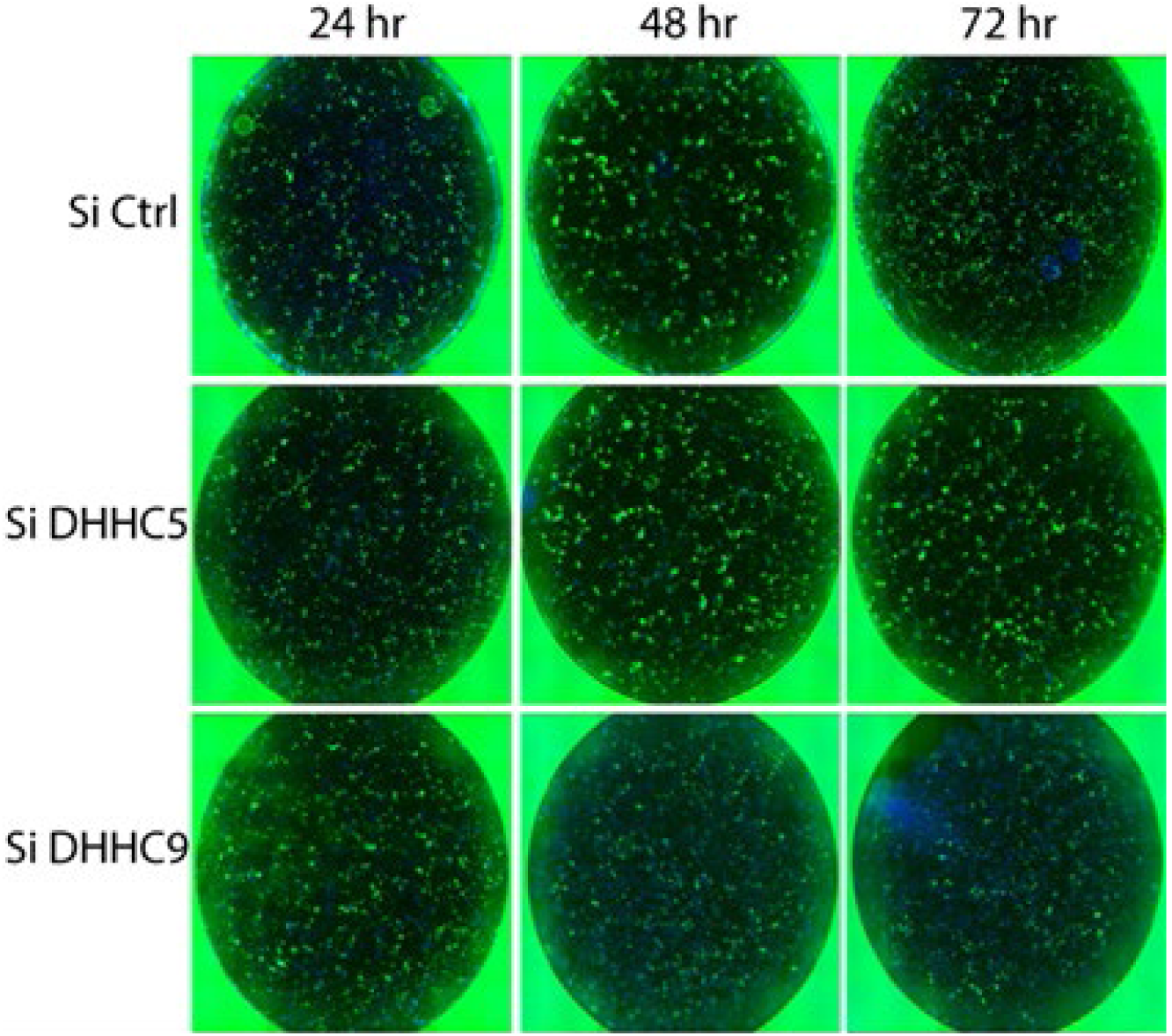
Supplemental image to Fig. 6A. Images of the entire well of the 96 well plate is provided to forego any bias in the region of interest photographed.

**Fig S4:**
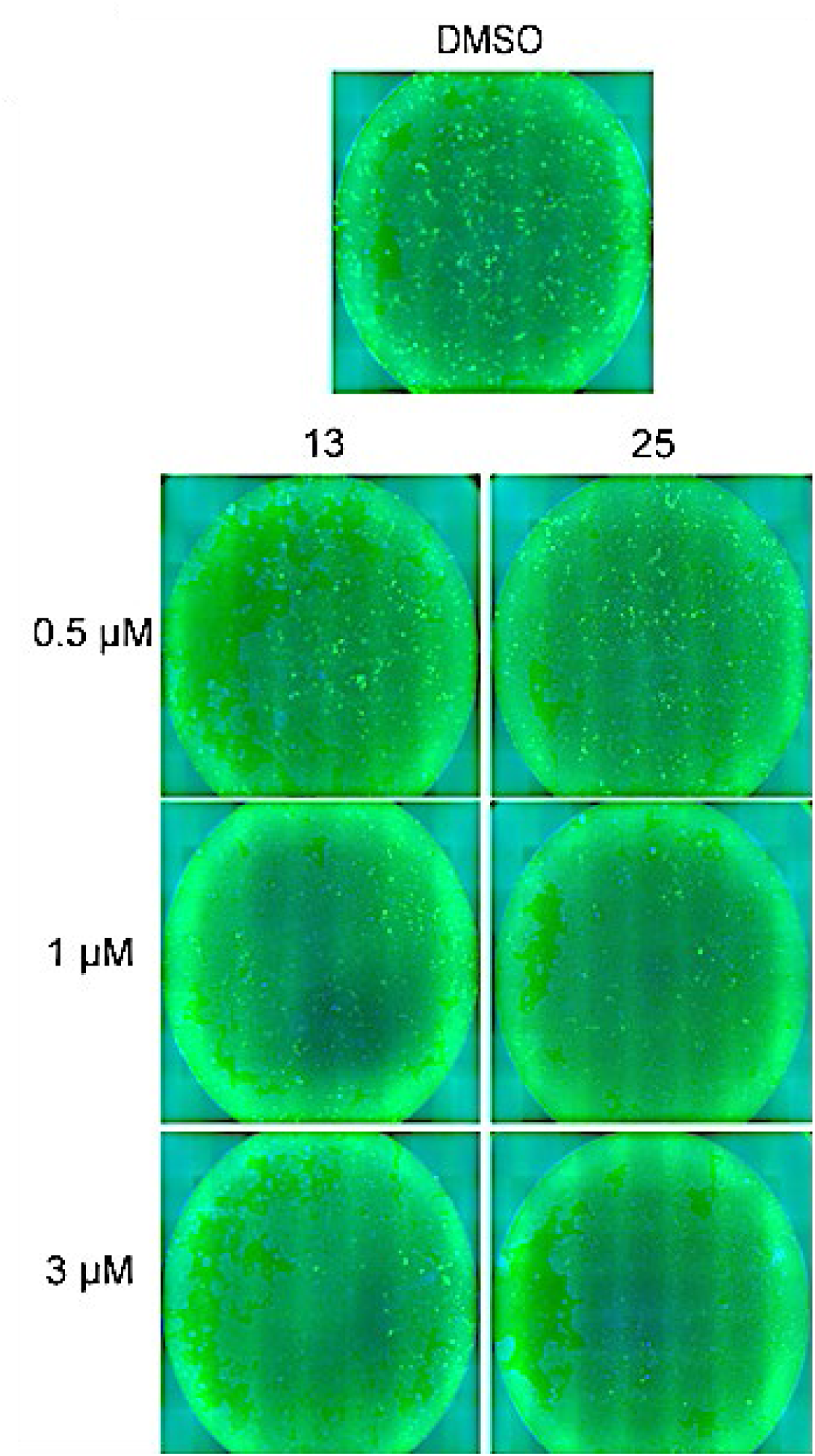
Supplemental image to Fig. 8A. Images of the entire well of the 96 well plate is provided to forego any bias in the region of interest photographed.

